# Active Surveillance Reveals a Systemic Pro-Resolving Th2 Immune Program Linked to Desmoid Tumor Regression

**DOI:** 10.64898/2026.04.16.718860

**Authors:** Laura Bergamaschi, Stefano Percio, Yiyi Zhu, Gabriele Tine, Rosalba Miceli, Marco Fiore, Elena Palassini, Paola Collini, Federica Perrone, Francesca Rini, Jessica Gliozzo, Cristina Banfi, Barbara Vergani, Biagio Eugenio Leone, Armando Giuseppe Licata, Loris De Cecco, Monica Zucchini, Arabella Mazzocchi, Sandro Pasquali, Alessandro Gronchi, Licia Rivoltini, Viviana Vallacchi, Chiara Colombo

**Author notes:** **Corresponding Author** Chiara Colombo Sarcoma Service Fondazione IRCSS Istituto Nazionale dei Tumori Milano Viviana Vallacchi Unit of Translational Immunology Department of Experimental Oncology Fondazione IRCSS Istituto Nazionale dei Tumori Milano. Equally contributing authors (Viviana Vallacchi, Chiara Colombo).

## Abstract

Desmoid fibromatosis (DF) is a rare mesenchymal neoplasm with an unpredictable clinical course, where spontaneous regression or progression occurs in a significant subset of patients through largely undefined mechanisms. The use of active surveillance (AS) offers the opportunity to investigate whether tumor- or host-driven systemic and local immune features may explain these divergent outcomes, improving patient management.

A prospective observational study enrolled 55 patients with primary sporadic DF managed with AS. Clinical evolution was categorized as progression, regression, or stable disease according to RECIST 1.1. Immunomonitoring with multicolor flow cytometry identified distinct systemic T-helper polarization states stratifying clinical trajectories: regressors showed a Th2-skewed profile, while progressors displayed activated T-helper cells and Th1/Th9/Th17 subsets. Higher baseline Th2 levels associated with regression and longer progression-free survival. Plasma proteomic and whole-blood transcriptomic analyses confirmed coordinated IL-4/IL-13–linked pro-resolving programs in regressors and inflammatory, early T-cell activation signatures in progressors. Tumor transcriptomics revealed adaptive, antigen-presentation and restrained immune programs in regressing lesions versus innate inflammatory, interferon and TGF-β–driven fibrotic pathways in progressing tumors.

These findings identify systemic T-helper polarization as a biomarker of DF behavior and highlight coordinated systemic–tumoral immune programs underlying clinical outcomes, supporting more precise clinical management.

## Introduction

Desmoid fibromatosis (DF) is a distinct, locally aggressive, mesenchymal neoplasm that affects young adults, with a female predominance. DF occurs either sporadically or in association with familial adenomatous polyposis (FAP) and is molecularly characterized by aberrant Wnt/β-catenin signaling, resulting from somatic *CTNNB1* exon 3 mutations, most frequently T41A and S45F, in sporadic form or germline APC alterations in FAP-associated cases ^1^.

Considering the unpredictable natural evolution, with nearly one third of patients experiencing spontaneous regression ^2–4^, active surveillance (AS) is recommended for the majority of patients ^5^. Spontaneous cancer regression, although generally uncommon, has long suggested that endogenous immune programs can restrain transformed cell growth ^6^. Classical paradigms of antitumor immunity posit that potent type-1 inflammation, characterized by Th1 polarization, cytotoxic T cell expansion, interferon-γ production, and M1 macrophage activation, underlies effective tumor control and may form the biological basis for therapies such as immune checkpoint blockade ^7,8^. However, this framework derives largely from established, immunogenic cancers and may not fully capture immune dynamics in pre-neoplastic or locally aggressive lesions. In contrast to chronic type-1 inflammation, which can perpetuate tissue damage and foster progression in early disease, specialized pro-resolving immune programs, including Th2-skewed responses, regulatory T cells, M2-like wound-healing macrophages, orchestrate the active resolution of inflammatory processes and restoration of tissue homeostasis and repair programs ^9–11^. Within this framework, DF, which shares features with wound-healing and fibroproliferative tissues, represents a unique model to investigate whether systemic and local type 2/reparative immune programs contribute to spontaneous regression, whereas persistent type 1-polarized inflammation may be associated with progression. This hypothesis is biologically plausible in light of the well-known role of homeostatic immune programs in remodeling the stromal environment and reprogramming local fibrosis ^12–15^, which is an intrinsic pathogenic process in DF^16^.

Since increasing evidence supports a contribution of the immune system to DF biology ^17–20^, we investigate whether clinical evolution of DF may be governed by distinct immune trajectories through the conduction of an integrated systemic and local immunoprofiling, with the final goal of defining predictive biomarkers of outcomes in patients with DF as well as novel therapeutic targets for this clinical setting.

## Results

### Patient Clinical Features

A total of 55 patients with histopathologic and molecular diagnosis of DF was enrolled. Patients entering the AS protocol underwent blood collection for longitudinal immunomonitoring, starting at baseline and then every 3 months during the first year and every 6 months during the second year of follow-up. Studies of the tumor microenvironment were performed on diagnostic FFPE biopsy specimens obtained at enrollment in AS program (**Fig. 1A and B**).

**Figure 1.**
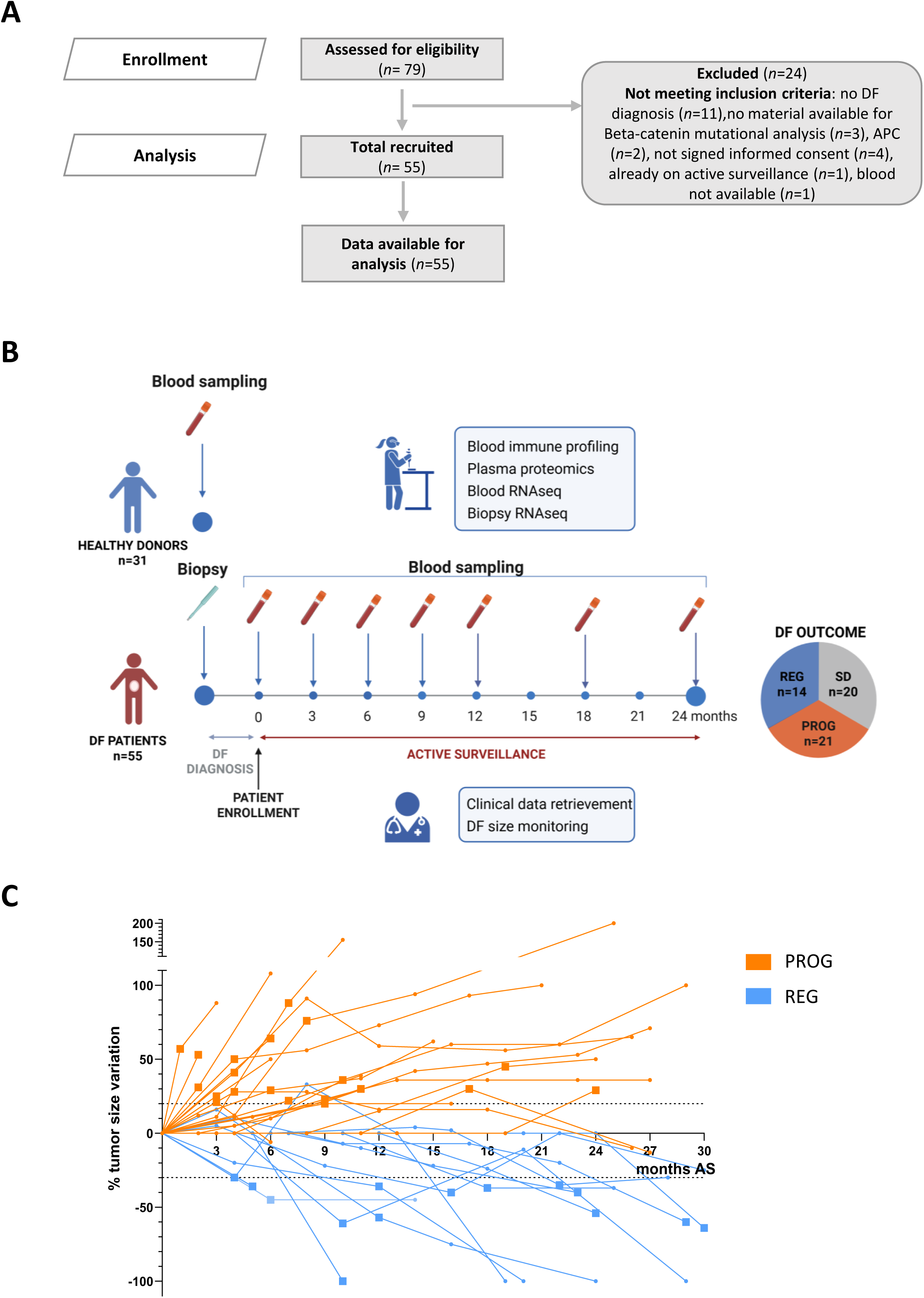
Study design. (A) Study diagram with exclusion criteria. (B) Graphical representation of the study design with DF patient’s outcome pie chart. (C) Spaghetti plot of DF patients AS, the RECIST events are represented with a squared timepoint. APC, adenomatous polyposis coli. AS, active surveillance. DF, desmoid fibromatosis. PROG, patients with RECIST progression as first event during AS. REG, patients with RECIST regression as first event during AS. SD, stable disease.

Forty-three patients were female (43/55, 78%), in line with DF epidemiological data ^21^. The vast majority of DF were located in the abdominal wall (29/55, 53%) and the most common *CTNNB1* mutation was T41A (28/55, 51%). Median tumor size was 50 mm (16-150 mm). The median follow-up was 48 months (IQR: 36-48 months). Over the AS period, 14 patients (25%) experienced disease regression (REG), 21 patients (38%) showed progression (PROG) as their first RECIST-event, and 20 patients (37%) had stable disease (SD). The median time to regression was 13.8 months (IQR: 7.0-22.2 months), while the median time to progression was 6.9 months (IQR: 4.1-10.4 months) (**Fig. 1C**), confirming the previous observation that regression tend to occur as a later event with respect to progression ^2^. A total of 18 patients (33%), 14 in the PROG group and 4 in the SD group, discontinued AS to initiate active treatment (32.7%), after a median time of 13 months (IQR 6-18) AS. During collection of anamnestic data, 11 of 52 patients (21%) reported a previous diagnosis of an autoimmune disease, including psoriasis, Hashimoto’s thyroiditis, systemic lupus erythematosus, Graves’ disease, vitiligo, celiac disease, scleroderma, and undifferentiated connective tissue disease (**Table 1, Suppl. Table S4**).

**Table 1.**
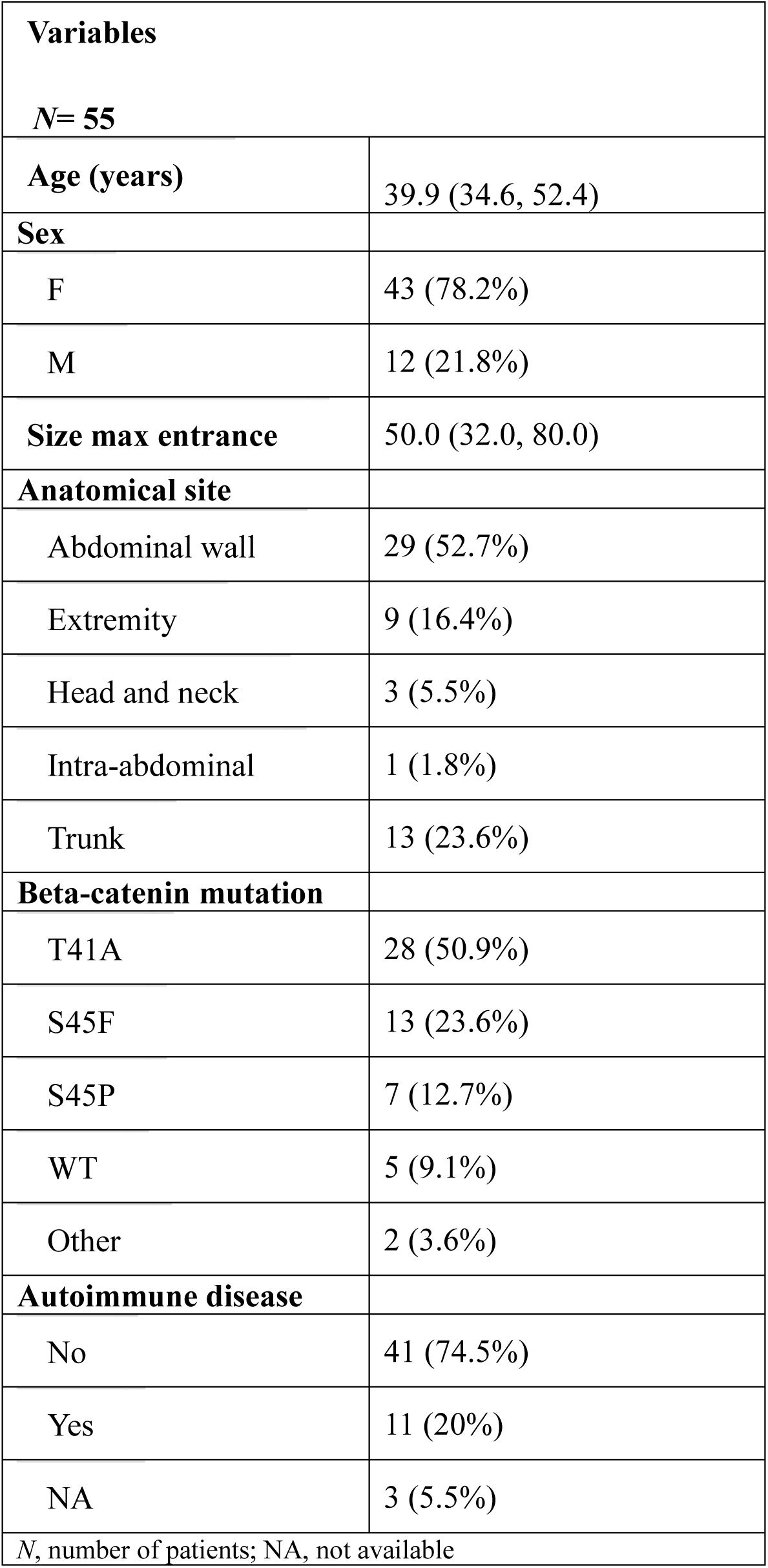
Patients’ Characteristics.

Kaplan-Meier curves were broadly overlapping for PFS across the predefined baseline clinical, pathological, and molecular categories (age, sex, *CTNNB1* mutation group, anatomic site group, and autoimmune disease status; **Suppl. Figure S7A-E**). Consistently, multivariate analysis for PFS showed no evidence of an independent association between baseline covariates and progression (**Table 2**), confirming the challenge in predicting outcomes of patients with DF. Only larger tumor size was independently associated with shorter TFS (80 vs 33 mm: HR = 4.44; 95% CI, 1.16-16.99; p = 0.003), while all other clinical-pathological and molecular features were not significantly associated with treatment initiation (age, p = 0.318; *CTNNB1* mutation status, p = 0.971; anatomic site, p = 0.529; and autoimmune disease, p = 0.267) (**Suppl. Figure S7E-L, Table 2**).

**Table 2.** Multivariate Cox model for progression-free survival and treatment-free survival.

| Variable | Progression-Free Survival |  |  | Treatment-Free Survival |  |  |
| --- | --- | --- | --- | --- | --- | --- |
|  | HR | 95% CI | p-value | HR | 95% CI | p-value |
| **Age\*** |  |  |  |  |  |  |
| 52 vs 35 years | 0.53 | 0.20, 1.43 | 0.264 | 0.82 | 0.28, 2.41 | 0.318 |
| **Beta-catenin mutation** |  |  |  |  |  |  |
| WT/S45P/Other | — | — |  | — | — |  |
| S45F | 0.47 | 0.12, 1.85 | 0.839 | 1.74 | 0.32, 9.55 | 0.971 |
| T41A | 0.41 | 0.11, 1.44 |  | 0.98 | 0.22, 4.42 |  |
| **Size\*** |  |  |  |  |  |  |
| 80 vs 33 mm | 1.13 | 0.40, 3.18 | 0.369 | 4.44 | 1.16, 16.99 | 0.003 |
| **Anatomical site** |  |  |  |  |  |  |
| Abdominal wall | — | — |  | — | — |  |
| Trunk/Ex/H&N/Intra-a | 3.18 | 0.95, 10.6 | 0.061 | 1.46 | 0.47, 4.56 | 0.529 |
| **Autoimmune disease** |  |  |  |  |  |  |
| No | — | — |  | — | — |  |
| Yes | 0.69 | 0.18, 2.59 | 0.584 | 2.08 | 0.60, 7.14 | 0.267 |

\* 52 and 32 years correspond to the 3rd and 1st quartile of age; 80 and 33 mm correspond to the 3rd and 1st quartile of tumor size.
List of abbreviations. HR, Hazard Ratio. CI, Confidence Interval. Wt, wild-type. Ex, Extremity. H&N, Head and Neck. Intra-a, Intra-abdominal.

#### Peripheral blood immune profiles at baseline distinguish DF patients from healthy donors but also serve as predictive markers for REG and PROG outcomes

DF carry a high risk of local recurrence after surgical intervention, likely due to tissue injury–induced activation of local myofibroblasts ^22^, underscoring the potential value of liquid biopsy–based biomarkers for disease monitoring. Leveraging on this clinical advantage, we investigated the distinctive characteristics of circulating immune cell populations in patients with DF under AS, searching for patterns associated with disease course and insight into the underlying immunological mechanisms. A multiparametric flow cytometry analysis, encompassing both myeloid and lymphoid cell populations, was performed in longitudinal samples prospectively obtained from fifty-two DF patients at AS program entering (baseline) and at different time points (3, 6, 9, 12, 18 months) during surveillance, including 20 PROG, 12 REG and 20 SD patients (**Suppl. Table S1, S4**). For comparison, blood samples from 31 age- and sex-matched healthy donors (HD) were prospectively collected and analyzed.

DF patients exhibited at baseline signs of dysfunctional immune cell profile when compared with HD, reflected by increased activated CD4⁺ T cells (CD3^+^CD4^+^HLA-DR^+^) and, within the CD4⁺ T-cell pool, a higher frequency of Th22 cells (defined as CD3^+^CD4^+^CCR4^+^CXCR3^-^CCR6^+^CCR10^+^). This was accompanied by an expanded myeloid compartment, characterized by increased total granulocytes, mature low-density granulocytes (CD15^+^CD11b^+^CD16^hi^), and non-classical monocytes (CD14^lo^CD16^+^). All these myeloid populations displayed also enhanced PD-L1 expression, pointing to systemic activation and regulatory polarization of both T and myeloid cells as a potential intrinsic feature of DF patients (**Fig. 2A**).

**Figure 2.**
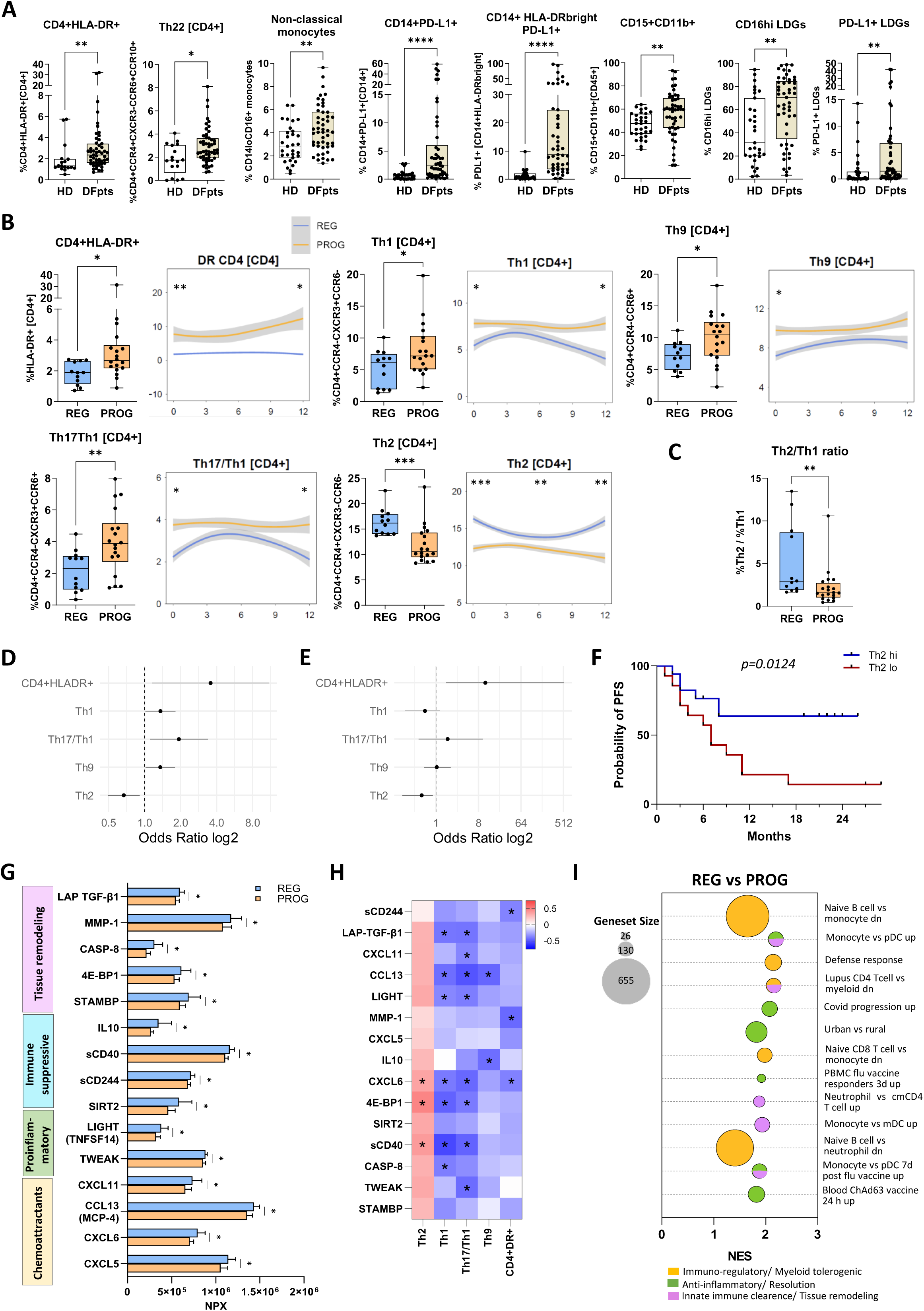
Peripheral blood immune characterization. (A) Box plots representing peripheral blood immune populations characterized by multiparametric flow cytometry in HD vs DF patients at diagnosis. (B) Box plots representing peripheral blood immune populations characterized by multiparametric flow cytometry in REG vs PROG DF patients at diagnosis. Regression plots of peripheral blood immune populations characterized by multiparametric flow cytometry in REG vs PROG DF patients during AS. (C) Box plot representing peripheral blood Th2/Th1 ratio in REG vs PROG DF patients at diagnosis. (D) Forest plot representing univariate logistic regression analysis of flow cytometry immune phenotyping data at diagnosis. (E) Forest plot representing multivariate logistic regression analysis of flow cytometry immune phenotyping data at diagnosis. (F) PFS curves of DF patients with high vs low percentages of Th2 at diagnosis, Long-rank test p=0.0124. (G) Bar chart representing plasma proteomics data obtained by Olink technology of REG vs PROG patients at diagnosis. (H) Heatmap representing linear correlations between peripheral blood immune populations and plasma protein levels. (I) Bubble plot representing REG vs PROG gene set enrichment analysis (GSEA) of peripheral blood transcriptional data tested for C7isdb, C7vax and custom immune signatures, categorized in Immuno-regulatory/Myeloid tolerogenic (yellow), Anti-inflammatory/Resolution (green), Innate immune clearance/Tissue remodeling (pink). AS, active surveillance. DF, desmoid fibromatosis. HD, healthy donors. LDG, low-density granulocyte. NES, normalized enrichment score. NPX, normalized protein expression. PFS, progression free survival. PROG, patients with RECIST progression as first event during AS. Pt, patient. REG, patients with RECIST regression as first event during AS. Th, T-helper cells. *p<0.05, **p<0.01, ***p<0.001 by Mann-Whitney test or Spearman correlation test.

To cluster baseline blood immune alterations associated with clinical outcomes in DF patients, we first compared baseline flow cytometry profiles between REG and PROG groups. A significant enrichment of Th2 cells (CD3⁺CD4⁺CCR4⁺CXCR3⁻CCR6⁻) within the CD3⁺CD4⁺ T-cell compartment was observed in REG patients. In contrast, PROG patients exhibited a markedly higher frequency of activated CD4⁺ T cells (CD3⁺CD4⁺HLA-DR⁺), along with increased proportions of Th1 (CD3⁺CD4⁺CCR4⁻CXCR3⁺CCR6⁻), Th9 (CD3⁺CD4⁺CCR4⁻CCR6⁺), and Th17/Th1 (CD3⁺CD4⁺CCR4⁻CXCR3⁺CCR6⁺) subsets. Notably, these differences, recapitulated also by the Th2/Th1 ratio (**Fig. 2C**), persisted throughout the first year of active surveillance in longitudinal blood samples (**Fig. 2B**), suggesting that the immune hallmarks possibly distinguishing REG with respect to PROG patients may represent a constitutive systemic immune background that might predispose to distinct disease behaviors. No significant differences were instead observed between groups in the myeloid compartment, regulatory T-cells, markers of T-cell exhaustion and immune checkpoint expression (**Suppl. Table S5**).

To increase the robustness of these findings, we conducted logistic regression analysis testing the association of these cell population abundance with the risk of first RECIST-event. Higher Th2 frequencies were associated with a statistically significant reduced risk of progression (OR = 0.67; 95% CI, 0.49-0.91) whereas increased activated CD4⁺ T cells, Th1, Th17/Th1 and Th9 CD4+ subsets were related with a statistically significant increased risk of progression (OR = 3.57, 1.36, 1.94 and 1.35, respectively) (**Fig. 2D, Suppl. Table S6**). When the presence of these CD4+ T-cells sub-populations were considered together in a multivariate analysis, patients with higher Th2 level at baseline were more likely to develop a tumor regression (OR = 0.50, CI 95%, 0.19-0.86; p = 0.046; **Fig. 2E**) independently from other sub-populations. Consistently, survival analysis showed that patients with high baseline Th2 levels (defined as above the median) had a longer PFS compared to those with low levels (log-rank test, p = 0.012; **Fig. 2F**).

For further increasing the robustness of these findings, we also considered patients with SD during AS and reclassified them based on any increase or decrease (even if not reaching RECIST threshold) (see Method section). The resulting groups, reported as rREG (N = 20) and rPROG (N = 32), respectively, confirmed a significantly higher frequency of Th2 CD4⁺ T cells in rREG patients and Th1/Th19/Th17 CD4⁺ T cells in rPROG patients (**Suppl. Figure S8**).

#### Integrated proteomic–transcriptomic profiling reveals distinct systemic immune programs in REG vs PROG outcomes

To gain functional insights into the systemic immune alterations captured by flow cytometry immunomonitoring, circulating cyto/chemokine profiling was performed by Olink platform alongside whole-blood transcriptomics in a selected subset of patients classified as PROG (N=16) or REG (N=10).

With respect to PROG, REG patients showed a baseline plasma immune proteomic milieu enriched in Th-2 CD4+ cells, revealed by the integration of *i*) chemokine-driven recruitment of reparative, alternatively activated myeloid and tissue-remodeling cells (CXCL11, CCL13/MCP-4, CXCL5, CXCL6); *ii*) concurrent regulatory and anti-inflammatory signals together with apoptosis-promoting factors (IL-10, soluble CD40, soluble CD244, SIRT2, LIGHT, TWEAK, CASP8); and *iii*) metabolic and translational control mechanisms favoring fibroblast clearance and extracellular matrix (ECM) degradation (LAP–TGF-β1, MMP-1, 4E-BP1, STAMBP) (**Fig. 2G, Suppl. Figure S9**). Collectively, this setting is compatible with the Th2-skewed, pro-resolutive immunity, in which regulatory cytokines might restrain chronic inflammation and metabolic brakes limit stromal proliferation, thereby favoring fibroblast clearance.

Consistent with this Th2-oriented immune context, the plasma factors IL-10, MMP-1, and TGF-β1, which were increased in REG, mapped to the Reactome “Interleukin-4 and Interleukin-13 signaling” pathway (**Suppl. Table S6**), a canonical pathway associated with Th2 polarization and type-2 immune responses. Circulating proteomic profile closely mirrored the flowcytometry-defined T-cell landscape, as CXCL6, 4E-BP1, and sCD40 correlated positively with Th2 frequencies, whereas LAP–TGF-β1, CCL13/MCP-4, LIGHT, CXCL6, 4E-BP1, sCD40, and CASP8 showed inverse associations with Th1 cells. Similar inverse relationships were observed for Th17/Th1, Th9, and activated CD4⁺HLA-DR⁺ T cells (**Fig. 2H**), indicating a coordinated regulation between circulating soluble mediators and disease-associated CD4⁺ T-cell phenotypes.

Peripheral blood transcriptomic profiles unraveled that REG patients were featured by a significant upregulation of immune-related signatures compared with PROG patients, consistent with both the flow cytometry and proteomic results (**Fig. 2I and Suppl. Table S8**). Indeed, anti-inflammatory and pro-resolutive pathways were marked by immuno-regulatory, tolerogenic myeloid signaling (*SIGLEC10*, *LILRB3*, *ZFP36L1*), anti-inflammatory mediators (*SERPINA1*, *SIRT2*), and metabolic restraint of oxidative stress and mTOR activity (*EIF4EBP2*, *SOD2*), alongside innate immune clearance and tissue-remodeling programs characterized by phagocytic and oxidative genes (*CD14*, *NCF2*), Fc/complement responses (*FCGR1A*, *C3AR1*), and antigen presentation machinery (*ITGAX*, *CD74*, *HLA-B*), consistent with controlled surveillance rather than overt inflammation (**Fig. 2I and Suppl. Table S8**). Analysis of patients using the defined rREG and rPROG groups, which incorporate RECIST-defined cases together with reclassified stable patients based on tumor dynamics during AS, confirmed all gene set differences, supporting the robustness of the systemic transcriptomic signatures associated with DF regression and progression. In addition, rPROG patients showed enrichment of gene sets related to early CD4⁺ T-cell activation and signal transduction, consistent with a state of increased T-cell activation (**Suppl. Table S9**).

#### DF biopsies displayed a consistently activated immune-infiltrated microenvironment

DF biopsies (N=34), including samples from 12 PROG, 8 REG, and 14 SD patients, unravel the consistent albeit variable presence of lymphocytic infiltration frequently organized into aggregates as evaluated by semi-quantitative analysis of diagnostic hematoxylin and eosin (H&E)–stained FFPE sections obtained at baseline. However, lymphoid infiltrate level did not apparently cluster with disease outcome in terms of REG vs PROG (**Suppl. Table S4; Fig. 3A and B**).

**Figure 3.**
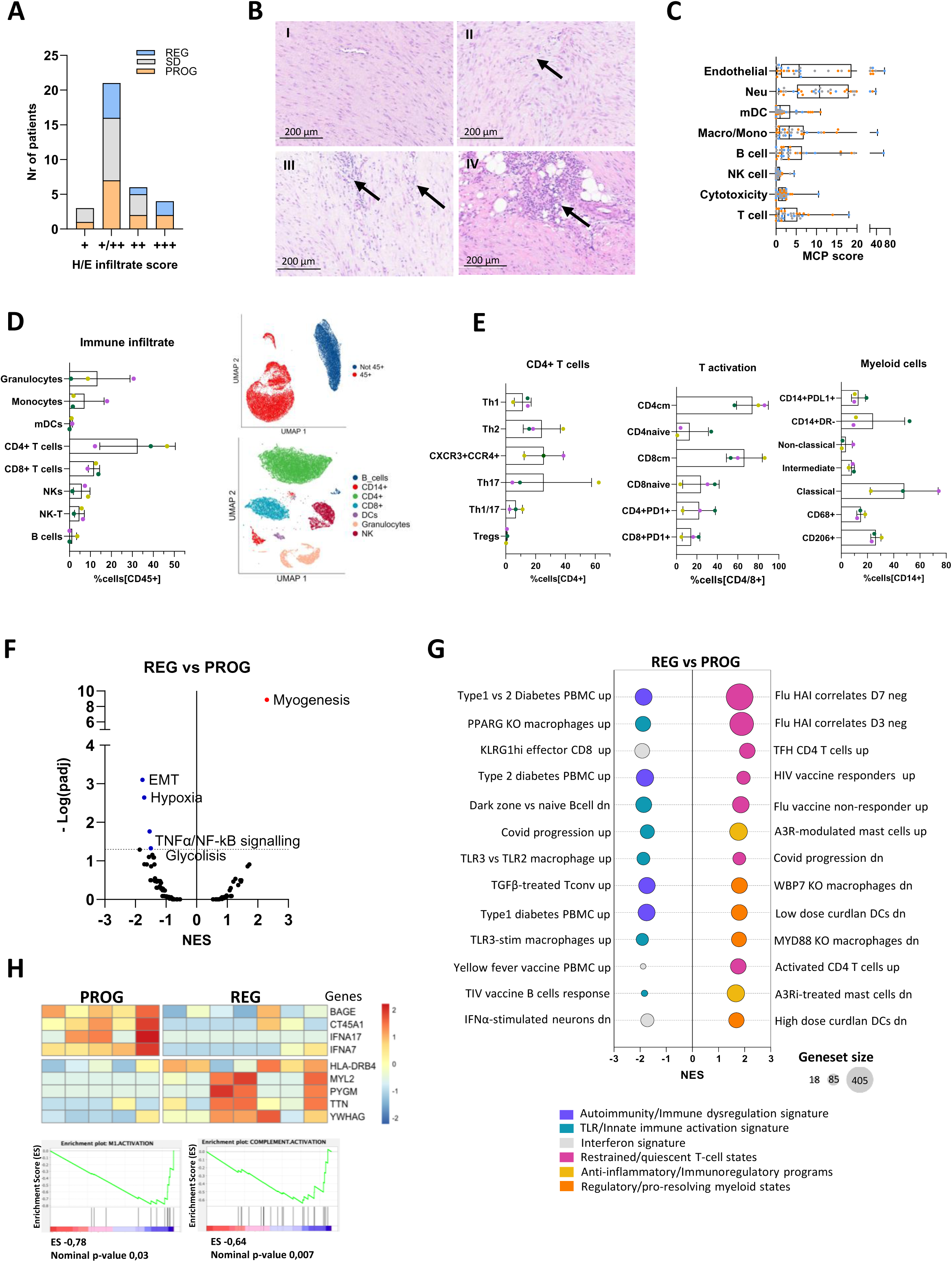
Tumor microenvironment characterization. (A) Bar chart representing semi-quantitative evaluation of lymphocytic infiltration on diagnostic H&E–stained FFPE sections from PROG, REG and SD patients categorized as weakly positive (+), moderately positive (+/++), positive (++) and strongly positive (+++). (B) Pictures of H&E-stained DF FFPE biopsies representative of weakly positive (I), moderately positive (II), positive (III) and strongly positive (IV) lymphocytic infiltrate, pointed by the black arrows. (C) Bar chart representing MCP-counter deconvolution analysis performed on DF FFPE biopsies RNAseq data. (D) Bar chart and UMAP plots representing the immune infiltrate present in DF surgical specimens characterized by multiparametric flow cytometry on single cell suspensions. (E) Bar chart representing further characterization of DF surgical specimens immune infiltrate by multiparametric flow cytometry. (F) Volcano plot representing GSEA of REG vs PROG biopsies transcriptional data, tested for Hallmark data set collection. (G) Bubble plot representing selected GSEA results of REG vs PROG biopsy transcriptional data tested for C7isdb, C7vax and custom immune signatures, categorized in Autoimmunity/Immune dysregulation signature (purple), TLR/Innate immune activation signature (green), Interferon signature (gray), Regulatory / pro-resolving myeloid states (orange), Anti-inflammatory / Immunoregulatory immune programs (yellow), Restrained / quiescent T-cell states (pink). (H) Heatmap and enrichment plots representing differentially expressed genes and signatures between REG and PROG identified by targeted transcriptional profiling of selected DF biopsies. EMT, epithelial to mesenchymal transition. FFPE, formalin fixed paraffine embedded. H/E hematoxylin and eosin. MCP, Microenvironment Cell Populations. mDC, myeloid dendritic cells. NES, normalized enrichment score. NK, natural killer. PROG, patients with RECIST progression as first event during AS. REG, patients with RECIST regression as first event during AS. SD, stable disease. Th, T-helper cell. TLR, tool like receptor.

To obtain integrated characterization of the tumor and immune microenvironment, RNA-seq experiment was performed on 34 DF biopsies (**Suppl. Table S4**). MCP-counter deconvolution method ^23^ indicated that major immune cell populations were detectable, including myeloid cells (monocytes/macrophages and dendritic cells), neutrophils, B cells, T cells, NK cells, and cytotoxic lymphocytes. Endothelial cell signatures were also consistently detected, reflecting the presence of a vascular compartment within the tumor microenvironment. Consistent with semiquantitative assessment of lymphoid infiltrates, no major differences in MCP-counter-derived immune cell abundance scores were observed between REG and PROG tumors (**Fig. 3C**).

To overcome the limitations of bulk RNA-seq in resolving the immune cellular composition and functional states within the DF tumor microenvironment, we performed multiparametric flow cytometry on single-cell suspensions obtained from surgical specimens of three DF patients.

Albeit referring to a very limited case set, due to the nowadays rare use of surgical maneuvers in DF patients ^5^, clinical data suggested that these specific cases could resemble PROG lesions. DF-derived cell suspensions showed the presence of CD45+ infiltrating immune cells, with a most prevalent CD4⁺ population (mean 32%) followed by granulocytes (mean 13%), CD8⁺ T cells (mean 11%) and CD14+ (mean 7%) monocytes. T cells displayed a central memory (CM) phenotype that reflected persistent antigen exposure and long-lived immune responses (**Fig. 3D and E**). Within the CD4⁺ T helper subsets, a concomitant presence of Th1, Th2, Th17, and Th1/17 cells expressing the chemokine receptors CXCR3 and CCR4, typically associated with Th1- and Th2-oriented migratory programs.

In contrast, regulatory T cells were scarcely detectable, suggesting limited immunoregulatory and homeostatic control (**Fig. 3D**). The myeloid compartment was also well represented and comprised classical and pro-inflammatory PD-L1 monocytes, neutrophils, myeloid dendritic cells (mDCs) and CD68+ or CD206+ macrophages. Similarly to Treg, immunosuppressive HLA-DR⁻ monocytes (M-MDSC) were also detected at low frequencies, further supporting the poorly immunoregulated nature of the DF milieu (**Fig. 3D and E**).

Collectively, these findings confirm the presence of robust immune system activation within DF lesions, displaying clear signs of immune activation. The concurrent presence of multiple helper and effector T-cell subsets, the paucity of regulatory mediators, and the abundance of inflammatory myeloid cells indicates uncoordinated and full-fledged immune response in a context of a chronic, non-resolving immune inflammation.

#### Pro-resolving versus myeloid–fibrotic programs may differentiate REG versus PROG DF transcriptomics

Enrichment analysis of DF biopsy transcriptomes (Hallmark, C7, and custom immune gene sets) identified distinct transcriptional programs in regressing (REG) versus progressing (PROG) tumors (**Suppl. Table S3**). REG tumors were enriched for the Myogenesis Hallmark gene set, indicating a more differentiated state, with leading-edge genes such as *MYH11* and *MEF2C*. In contrast, PROG tumors showed enrichment of disease-promoting gene sets including Epithelial–Mesenchymal Transition, Hypoxia, TNF-α/NF-κB signaling, and Glycolysis, reflecting inflammatory activation, metabolic rewiring, and fibrotic stromal remodeling. Their leading-edge genes included ECM- and hypoxia-related factors (*LOXL2*, *MMP14*, *VEGFA*) and inflammatory mediators (*CXCL11*, *NR4A1*) (**Fig. 3F, Suppl. Table S10**). Immune gene set analysis further highlighted divergent immune landscapes (**Fig. 3G and Suppl. Table S10**). REG tumors exhibited coordinated adaptive immune responses, with enrichment of restrained/quiescent T-cell states (e.g., *IL7R*), antigen presentation (MHC class II), dendritic-cell differentiation, and immunoregulatory circuits, consistent with a controlled, memory-oriented immune microenvironment. In contrast, PROG tumors displayed limited adaptive engagement, restricted to transient early T-cell activation (NR4A genes), alongside innate inflammatory signatures, including TLR activation, type I interferon responses, immune dysregulation modules, neutrophil-associated genes (*FCGR3B*, *S100A8*/*A9*), and TGF-β–driven fibrogenic remodeling. Reclassification analysis confirmed these differences (**Suppl. Table S11**).

Overall, regressing DF tumors are characterized by differentiated tumor programs and regulated adaptive immunity, whereas progressing tumors show inflammatory, innate-dominated, and fibrotic transcriptional profiles associated with adverse clinical outcomes.

To further validate the RNA-seq findings using an FFPE-compatible approach, we analyzed a subset of DF samples (7 REG and 5 PROG) applying a targeted platform interrogating immune and tumor-related gene panels (**Suppl. Table S4**). This analysis largely recapitulated the transcriptional differences observed by RNA-seq, with REG tumors retaining features of fibroblastic differentiation, whereas PROG lesions showed enrichment of inflammatory immune programs, including M1-associated activation and complement-related pathways (**Fig. 3H**). Overall, these results confirm the robustness of the REG- versus PROG-associated transcriptional signatures across independent technologies and sample-processing workflows.

#### Autoimmune disease is associated with enhanced intratumoral immune activation in DF

The above described targeted transcriptomic analysis revealed the expression of HLA-DRB4 in 5 of 12 DF biopsies (42%) (**Fig. 3H**), a frequency higher than that reported in the general Western population (∼28%); ^24^. The comparison with frequencies reported in the Western population was performed using DRB4*01 positivity, which in our cohort corresponds to the presence of the DRB4 gene. Because of the anamnestic data on the high frequency of autoimmune disease in our DF population, we further explored the potential relationship between HLA-DRB4, autoimmunity, and DF clinical behavior, by first assessing HLA-DRB genotype frequency in 52 DF patients (**Suppl. Table S4**). The HLA-DRB4 gene, present in subjects carrying either HLA-DRB1*04, HLA-DRB1*07, or HLA-DRB1*09 allele groups, was identified in 22 of 52 patients (42%) (**Suppl. Table S12**). HLA-DRB4 was detected in 8 of 13 REG patients (62%) and in 10 of 21 PROG patients (48%), but this difference was not statistically significant. Among the 52 patients with available HLA-DRB4 data, autoimmune history was known for 49 (11 with autoimmune disease and 38 without). HLA-DRB4 was present in 7 of 11 patients with autoimmune disease (64%) compared with 15 of 38 patients without autoimmunity (39%). This 25% difference did not translate into statistical significance (Fisher’s exact test, p=0.180) likely due to small sample size.

Differential peripheral immune profiling was then analyzed to further investigate whether previous autoimmune disease could impact on DF systemic immunity. A largely comparable blood immune composition both in terms of cell populations and the soluble factors was observed in patients with and without autoimmune disease (**Fig. 4A**). Plasma factors found to be significantly increased in patients with autoimmune disease were sCD6, CDCP1, CCL4, IL-6, and IL-8, representing markers of generalized chronic immune activation commonly observed across heterogeneous autoimmune disorders (**Fig. 4B**) but apparently playing no role in differentiating our REG vs PROG DF patients. Conversely, tumor-level analyses identified marked differences in the tumor microenvironment, with multiple transcriptional signatures related to immune activation remarkably increased in DF lesions from patients with autoimmune disorders (N = 8) compared with those without (N = 25) (**Fig. 4C and D, Suppl. Table S13**).

**Figure 4.**
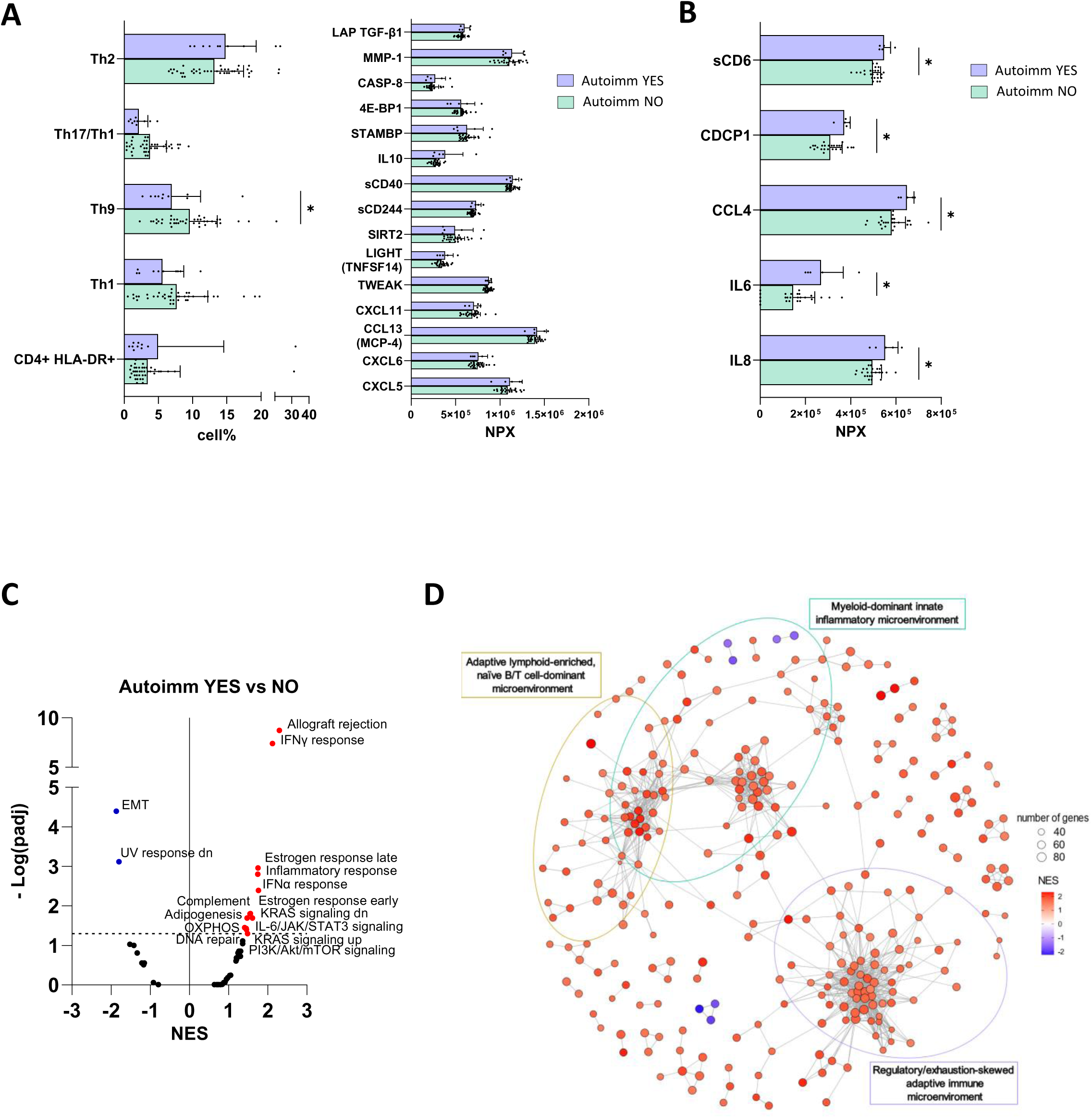
Characterization of the influence of autoimmune diseases in DF patients peripherasl blood and TME. (A) Bar chart representing selected peripheral blood immune populations and plasma proteomics data, obtained by multiparametric flowcytometry and Olink technology respectively, of Autoimm YES vs NO patients at diagnosis. (B) Bar chart representing significant plasma proteomics data obtained by Olink technology of Autoimm YES vs NO patients at diagnosis (C) Volcano plot representing GSEA of Autoimm YES vs NO biopsy transcriptional data, interrogated with Hallmarks data set collection. (D) Enrichment map representing the network of significantly enriched pathways identified by GSEA (p_adj_<0.05) of Autoimmunity YES vs NO biopsy transcriptional data tested for C7isdb gene set collection. Edges represent Jaccard similarity of overlapping leading-edge genes (edge weight > 0.1) and the ellipses highlight the top-3 clusters by number of pathways. Autoimm NO, DF patients without autoimmune diseases. Autoimm YES, DF patients with autoimmune diseases. NES, normalized enrichment score. *p<0.05 by Mann-Whitney test.

DF tumors from patients with autoimmune disease showed positive enrichment of immune-related pathways (IFN-γ/IFN-α, inflammatory signaling, complement, IL6–JAK–STAT3), together with oxidative phosphorylation, and DNA repair programs, and negative enrichment of stromal/mesenchymal gene sets (EMT, TGF-β–driven ECM remodeling) (**Fig. 4C**, **Suppl. Table S13**). Consistent with these findings, GSEA of C7 immunologic signatures indicated a coordinated adaptive immune landscape largely overlapping with the “Adaptive lymphoid–enriched, naïve B/T cell–dominant microenvironment”, characterized by enhanced antigen-presentation programs with increased MHC class I/II expression and enrichment of B-cell (e.g., *CD19*, *MS4A1*) and T-cell priming/memory (e.g., *IL7R*, *CCR7*) gene signatures. Inflammatory and myeloid-associated modules (e.g., *FGL2*, *PLBD1*) were also enriched but accompanied by regulatory and negative-feedback pathways, indicating a controlled activation state with similarities to both the “Regulatory/exhaustion-skewed adaptive immune microenvironment” (e.g., *TRAFD1, IRF1*) and the “Myeloid-dominant innate inflammatory microenvironment”. Stromal and ECM-remodeling gene signatures (e.g. *LOXL2*, *THBS2*) were reduced (**Fig. 4D, Suppl. Table S13**).

Overall, autoimmune-associated DF lesions display an immune-dominant, coordinated adaptive landscape with enhanced antigen presentation.

## Discussion

In this prospective clinical study, peripheral immune profiling revealed that patients with DF under AS display broad systemic immune perturbations at diagnosis compared with healthy donors, involving both lymphoid and myeloid compartments and consistent with a state of global immune dysregulation. Although DF is clinically characterized by local aggressiveness, the systemic immune alterations observed here may reflect the combined effects of tumor-driven inflammation and host-specific immunophenotypes. Actually, the balance and the dynamic interplay between tumor- and host-related factors appears to shape the systemic immune contexture, potentially influencing disease course. Indeed, patients who subsequently underwent spontaneous regression exhibited increased circulating Th2 cells and a coordinated peripheral immune program consistent with a pro-resolving and anti-inflammatory phenotype, characterized by immunoregulatory chemokines, apoptotic pathways, metabolic restraint and regulated innate immunity. In contrast, patients developing DF progression displayed expansion of activated Th1, Th9 and Th17/Th1 subsets, together with systemic signatures indicative of heightened T-cell activation in the absence of effective regulatory balance, consistent with a more inflammatory and less resolved immune state. The systemic immune profile observed in DF appears to align more closely with tissue repair and pro-resolving stromal–immune programs than with the immune-mediated mechanisms reported in other spontaneously regressing contexts, such as neuroblastoma, where lymphocyte activation and immune surveillance have been implicated, or nodular fasciitis, whose self-limited behavior has been linked to interferon-associated signaling pathways ^25,26^. The evidence that the differences in circulating T-helper balance persisted during the first year of AS is in line with the hypothesis that Th polarization is influenced by individual intrinsic immunotypes ^27^. If validated in a larger clinical cohort, the Th2/Th1 blood ratio could emerge as a promising biomarker for early risk stratification, leveraging a simple liquid biopsy in a tumor where surgical intervention may heighten the risk of recurrence. Therapies such as IL-4/IL-13 agonists or low-dose IL-2 variants, which promote Th2 differentiation in autoimmunity or allergy ^28^, could also in the future be repurposed in DF to promote immune rebalancing and possibly sustain regression even in progressing lesions. Notably, some current DF treatments, including NSAIDs, TKIs and low dose chemotherapy ^5^, may already bias the immune response toward Th2 in vivo, supporting the concept that Th2 polarization itself may represent a novel, targetable axis for more specific and personalized DF therapy.

Th2-polarized immunity has traditionally been associated with pro-tumorigenic functions, largely due to its ability to dampen Th1-driven cytotoxic responses, promote immunoregulatory myeloid compartments, and support stromal remodeling to allow cancer progression ^29^. Nevertheless, this paradigm is increasingly being reconsidered in a context-dependent manner. Emerging evidence indicates that Th2 responses may also exert anti-tumor or disease-limiting effects depending on tissue architecture, immune composition, and disease stage ^30–32^. In precancer lesions, recent data revealed that Th2 immunity may directly trigger a local cascade that clears premalignant cells ^32^, thus attributing to anti-inflammatory immune response a previously unrecognized role in counteracting cancer initiation ^33^.

Th2-associated programs have been also implicated in the resolution of inflammation and in tissue repair processes, acting to limit prolonged inflammatory stress and promote restoration of tissue homeostasis ^34^. This role is well established in skeletal muscle injury, where Th2 cells induce IL-4–dependent myeloid cell polarization toward a restorative phenotype, with macrophages removing necrotic debris, stimulating angiogenesis, and enhancing muscle progenitor activation, thereby driving tissue recovery. Consistently, Th2 cells are indispensable for muscle healing, and higher circulating Th2 levels associate with optimal regenerative outcomes ^35–38^. In the context of DF, a key unresolved question concerns the nature of the tissue perturbation that may initiate this reparative program. Given that the majority of DF in our cohort carried activating *CTNNB1* mutations (**Table 1**), β-catenin activation may establish a common fibroproliferative, wound-like state, while the clinical evolution of this state may depend on the immune context and, at least in part, on mutation-specific tumor biology, as also suggested by the less favorable outcomes reported for patients with S45F-mutant tumors during AS ^39^. The evidence that persistent Th1/M1 inflammatory environment is conversely linked with prolonged inflammation and dysfunctional repair, further suggests that analogous Th/macrophage immunoregulatory dynamics may shape whether the lesion is contained, remodeled, or driven toward fibroproliferative expansion in DF patients.

Consistently with the remarkable systemic alterations, also diagnostic DF biopsies exhibited in our case set a highly immune-infiltrated tumor microenvironment, characterized by a broad and heterogeneous immune cell composition, including myeloid cells, neutrophils, T and B lymphocytes and NK cells, in line with previous published reports ^17,20,40^. Transcriptional DF profiling revealed that progressive tumors are characterized by a coordinated inflammatory–fibrogenic program, integrating hypoxia adaptation, type I interferon signaling, neutrophil-driven myeloid inflammation, TGF-β–dependent stromal remodeling with T-cell activation. This inflammatory–fibrogenic state contrasts with regressive DF, which maintains a more differentiated fibroblastic identity and exhibits a pro-resolving, adaptive immune ecosystem enriched for memory-biased T-cell programs, and a basal interferon-driven program that supports immune readiness without triggering overt inflammatory responses. Together, these observations suggest that progressive and regressive DF lesions may follow distinct inflammatory–repair trajectories already established at diagnosis.

The prevalence of autoimmune diseases in our DF cohort was higher than that reported in the general Italian population, where autoimmune conditions affect approximately 1-10% of individuals depending on the specific disease ^41,42^. Autoimmune disorders are increasingly recognized as clinical manifestations of chronic immune dysregulation, and in the context of solid tumors they have been associated with context-dependent effects on tumor development and immune-mediated control ^43,44^. Within this framework, the HLA-DRB4 gene has previously been linked to autoimmune conditions and states of heightened or deregulated immune responsiveness ^45–48^. Together, these findings provided the rationale to investigate whether autoimmune comorbidities are associated with distinct immune features in DF that could impact outcome. Results showed that DF lesions arising in the context of autoimmune disease display a highly immune-engaged tumor microenvironment. However, neither the presence of autoimmune disease nor carriage of the HLA-DRB4 gene was associated with disease evolution, suggesting that autoimmune-related immune alterations may shape DF susceptibility and immune contexture, but are insufficient to directly determine clinical trajectory under AS. Notably, no major differences in systemic immune composition were observed between patients with and without autoimmune disease, further supporting the notion that the systemic immune signatures distinguishing progressive from regressive DF reflect intrinsic immune predispositions of the host, rather than secondary immune perturbations driven by autoimmune comorbidity.

Our study has several limitations. The relatively small sample size, particularly when patients were stratified according to clinical outcome, inevitably limited statistical power although this was partially mitigated by reclassifying stable patients based on longitudinal tumor size dynamics during AS. In addition, the single-center design and the rarity of DF may limit the generalizability of our findings, warranting validation in larger multicenter prospective cohorts, which is presently ongoing (NCT07496242). Longitudinal immune profiling beyond the first year of active surveillance was not feasible, as most patients experiencing progression during AS initiated treatment shortly after progression, precluding direct comparison of long-term systemic immune trajectories. Furthermore, tumor immune analyses were restricted to diagnostic biopsy specimens, limiting tissue availability and preventing spatially resolved and longitudinal assessment of immune–stromal interactions during disease evolution.

In summary, our findings support a model in which progressive and regressive DF lesions represent divergent inflammatory–repair trajectories detectable both systemically and at the tumor level. In this context, host immune predisposition, particularly the Th2/Th1 balance, may influence whether immune-regulatory mechanisms required for resolution are engaged. A Th2-biased systemic profile may favor immune regulation and tissue repair, whereas a Th1-skewed state may sustain unresolved inflammation and promote continued tumor progression under AS. These findings highlight the potential clinical utility of blood-based immune biomarkers, for risk stratification at diagnosis and informed patient management under AS. The hypothesis that immune-modulating strategies aimed at rebalancing immune polarization toward type-2–associated, pro-resolutive programs could be also explored for progressive DF.

## Methods

### Patients cohort

Patients with primary sporadic DF presenting with measurable disease in any anatomical site were prospectively enrolled in an observational study on active surveillance (AS) at our Institution between April 2019 and September 2022 (**Fig. 1A**). The study protocol and amendments were approved by Institutional Review Boards in accordance with applicable regulations (INT 90/18). This study was conducted in compliance with the principles outlined in the Declaration of Helsinki. Written informed consent was obtained from all participants prior to study inclusion.

Eligible patients were required to be at least 18 years old and have a biopsy-proven, histologically and molecularly confirmed diagnosis of DF located in the head and neck, trunk, abdominal wall, intra-abdominal cavity, or extremities. Patients with DF harboring *APC* gene mutations, recurrent disease, or prior primary surgical resection were excluded from the study. *CTNNB1* mutational status was assessed using direct Sanger sequencing as previously described ^49^. If *CTNNB1* appeared wild-type by Sanger sequencing, genomic sequencing was performed using the Ion Torrent platform with a targeted panel, the Hot-spot Cancer Panel (Ion AmpliSeq Cancer Hotspot Panel v2 #4475346, Thermo Fisher Scientific, Waltham, MA, USA).

Tumor size was evaluated considering the largest diameter of DF on contrast-enhanced magnetic resonance imaging (MRI) or computed tomography (CT) scans, with CT imaging primarily utilized for intra-abdominal lesions or when MRI was not feasible. Tumor measurements were recorded at diagnosis and throughout active surveillance, with follow-up imaging performed every three months during the first year and every six months during the second year, or until the start of active treatment for disease progression.

Core needle biopsy of the DF lesion was obtained at diagnosis and blood samples were collected both at baseline and at predefined time points during AS (3, 6, 9, 12, 18 and 24 months, or until transition to any active treatment) (**Fig 1B**). Variations on tumor largest diameter during AS was assessed according to RECIST version 1.1 ^50^. Tumor progression was defined as a > 20% increase in the largest diameter from baseline, partial regression as a > 30% reduction, and stable disease as any variation failing to meet either threshold. Patients were classified as having progression (PROG) or regression (REG) based on the first RECIST-defined event observed during their disease course. To improve the robustness of outcome-associated immune analyses, patients with RECIST-defined stable disease, who are not captured by the initial classification, were further stratified according to the direction of tumor size change during AS. Specifically, stable patients showing a progressive decrease in the largest tumor diameter over time were reclassified as regressors, whereas those showing a progressive increase were reclassified as progressors. These reclassified stable cases were then integrated with RECIST-defined PROG and REG patients to generate the reclassified groups rREG and rPROG, including 20 and 32 patients, respectively.

The decision to initiate active treatment upon DF progression defined as radiological and/or clinical progression was finally made at the clinician’s discretion in consultation with the patient. Treatment regimens were individualized and included locoregional therapies such as cryotherapy and electrochemotherapy, as well as systemic therapies including hormonal agents (e.g., tamoxifen, toremifene) and low-dose chemotherapy (e.g., methotrexate and vinorelbine).

### Blood collection, plasma and PBMC isolation

Peripheral blood samples from DF patients were collected at diagnosis and at the above-mentioned predefined time points during AS. Samples from age- and sex-matched healthy donors were also collected. Blood draws were performed in 10 mL Vacutainer tubes containing spray-coated K₂EDTA (BD Vacutainer™ Hemogard™ Closure Plastic K2-Edta Tube, #367525, Becton Dickinson, Franklin Lakes, NJ, USA). A 100 µL aliquot of blood was immediately analyzed by multicolor flow cytometry. Plasma was separated by two consecutive centrifugations at 1300 × g and stored at −80°C until further use. Peripheral blood mononuclear cells (PBMCs) were isolated from the remaining cellular fraction using Ficoll density gradient centrifugation (Greiner Bio-One Leucosep Polypropylene Centrifugation Tubes, #10030322, Thermo Fisher Scientific, Waltham, MA, USA) at 800 × g for 15 minutes without braking. The PBMC layer was carefully collected, washed with PBS, and residual red blood cells were lysed using Lysing Buffer (BD Pharm Lyse™ Lysing Buffer, # 555899, Becton Dickinson, Franklin Lakes, NJ, USA). Cell count was determined using a Bürker counting chamber, and viability was assessed with Trypan blue dye exclusion (Trypan Blue solution, #T8154-100ML, Sigma-Aldrich, St. Louis, MO, USA). One million PBMCs per staining condition were immediately analyzed by multicolor flow cytometry, while the remaining cells were cryopreserved in RPMI medium supplemented with 30% human serum and 10% dimethyl sulfoxide (Dimethyl Sulfoxide, #D2650-100ML, Sigma-Aldrich, St. Louis, MO, USA) and stored in liquid nitrogen for subsequent analyses.

### Dissociation of fresh DF surgical specimens

DF surgical specimens cell suspensions were obtained to perform multiparametric flow cytometry. After a first mechanical tissue dissociation, an enzymatic digestion was performed using Miltenyi Tumor Dissociation kit (Tumor Dissociation Kit, human, #130-095-929, Miltenyi Biotec, Bergisch Gladbach, Germany), GentleMACS Dissociator (gentleMACS™ Dissociator, #130-093-235, Miltenyi Biotec, Gladbach, Germany) and MACS Tube Rotator (MACSmix™ Tube Rotator, # 130-090-753, Miltenyi Biotec, Gladbach, Germany), following the manufacturer instructions for “Tough” and 0.2-1 g samples. The obtained suspension was 70 µm filtered (ClearLine® cell strainers 70 um, Biosigma S.p.a., Cona, Italy), washed with PBS and residual red blood cells were lysed using Lysing Buffer (BD Pharm Lyse™ Lysing Buffer, # 555899, Becton Dickinson, Franklin Lakes, NJ, USA). Cell count was determined using a Bürker counting chamber, and viability was assessed with Trypan blue dye exclusion (Trypan Blue solution, #T8154-100ML, Sigma-Aldrich, St. Louis, MO, USA). The obtained cells were immediately analyzed by multicolor flow cytometry.

### Flow cytometry

Immunophenotyping was performed by multicolor flow cytometry on whole blood, fresh or cryopreserved PBMCs, and fresh or cryopreserved single-cell suspensions derived from surgical specimens. The antibody panels used to evaluate myeloid and lymphoid immune populations are detailed in **Suppl. Table S1**. Briefly, samples were pre-treated with an FcR blocking reagent (FcR Blocking Reagent, human, #130-059-901, Miltenyi Biotec, Gladbach, Germany) and, when required, incubated with a live/dead viability dye (LIVE/DEAD™ Fixable Violet Dead Cell Stain Kit, for 405 nm excitation # L34963; LIVE/DEAD™ Fixable Near IR (876) Viability Kit, for 808 nm excitation #L34980, Thermo Fisher Scientific, Waltham, MA, USA). Cells were then stained with various conjugated monoclonal antibodies (mAbs) listed in **Suppl. Table S2**. Following staining, samples were washed and fixed in PBS containing 1% formalin. For intracellular staining, fixation and permeabilization were performed using an intracellular fixation and permeabilization buffer set (eBioscience™ Intracellular Fixation & Permeabilization Buffer Set, # 88-8824-00, Thermo Fisher Scientific, Waltham, MA, USA). Sample acquisition was carried out using a CytoFLEX S flow cytometer (CytoFLEX S, Beckman Coulter, Brea, CA, USA). Data were analyzed with Kaluza Analysis Software (Kaluza Analysis Software, version 2.1, Beckman Coulter, Brea, CA, USA) and expressed as the percentage of positive cells, with marker thresholds determined based on the corresponding isotype control. The applied gating strategies are reported in **Suppl. Figure S1-S6**. Data visualization, exploration, and statistical analysis were conducted using GraphPad Prism (GraphPad Prism, version 10, Dotmatics, Boston, MA, USA) or R software (R software, version 4.4.1, R Foundation for Statistical Computing, Vienna, Austria).

### Circulating proteome profiling

Plasma samples were analyzed using the Olink Target 96 Inflammation panel (Olink Target 96 Inflammation, Thermo Fisher Scientific, Waltham, MA, USA), which employs Proximity Extension Assay (PEA) technology. This method combines an antibody-based immunoassay with the high sensitivity of real-time polymerase chain reaction (PCR) ^51^. Threshold cycle (Ct) values obtained from internal and external controls underwent quality control and normalization. Protein levels were quantified on a relative scale and reported as normalized protein expression (NPX) units on a log₂ scale, where higher NPX values indicate higher protein concentrations. NPX values were IPC-normalized. Data visualization, exploration, and statistical analyses were performed using GraphPad Prism (GraphPad Prism, version 10, Dotmatics, Boston, MA, USA) or R software (R software, version 4.4.1, R Foundation for Statistical Computing, Vienna, Austria).

### RNA extraction from whole blood and transcriptional profiling analysis

Whole blood RNA was extracted from 5 ml of blood collected in PAXgene blood RNA tubes (PAXgene Blood RNA Tubes (IVD), #762165, PreAnalitiX GmbH, Hombrechtikon Switzerland) using the Blood miRNA Kit (PAXgene Blood miRNA Kit, #763134, PreAnalitiX GmbH, Hombrechtikon Switzerland), following the manufacturer instructions. The extracted RNA was quantified by NanoDrop 2000 spectrometer (NanoDrop™ 2000 Spectrophotometers, #ND-2000, Thermo Fisher Scientific, Waltham, MA, USA). The RNA yield ranged from 30 to 275 ng/µl. RNA quantity and quality were evaluated using Qubit RNA HS Assay Kit (Qubit RNA HS Assay Kit, #Q32852, Thermo Fisher Scientific, Waltham, MA, USA) by Qubit fluorometer (Qubit 4 fluorometer, Thermo Fisher Scientific, Waltham, MA, USA) and Bioanalyzer (2100 Bioanalyzer Instrument, Agilent, Santa Clara, CA, USA). RNA integrity number (RIN) values ranged from 7.6 to 9.5, while DV200 values varied from 77% to 99%. RNA-seq libraries were generated from 0.2 μg of RNA using Illumina Total RNA Prep with Ribo-Zero Plus (Illumina® Stranded Total RNA Prep, Ligation with Ribo-Zero Plus (96 Samples), # 20040529, Illumina Inc, San Diego, CA, USA) according to manufacturer’s recommendations. Libraries were sequenced on Illumina NovaSeq6000 (NovaSeq6000, Illumina Inc, San Diego, CA, USA) in 2X100-nt-long paired-end read modality. FastQC method was perform a quality check of FASTQ files ^52^. Fastp and SortMeRNA, and STAR algorithms were used to preprocessed and align reads to reference genome GRCh38, respectively ^52–54^. RSEM method was used to generate raw count matrix with hs_ensembl.h38.v112 annotation parameter ^55^.

### RNA extraction from DF FFPE biopsies and transcriptional profiling

RNA was extracted from formalin-fixed, paraffin-embedded (FFPE) DF biopsy sections using the miRNeasy FFPE Kit and the automated QIAcube extractor (QIAcube Connect, #9002864, Qiagen, Hilden, Germany), following the manufacturer’s instructions. RNA quantification was performed using the RNA HS Assay Kit (Qubit RNA HS Assay Kit, #Q32852, Thermo Fisher Scientific, Waltham, MA, USA) and a Qubit 4 Fluorimeter (Qubit 4 fluorometer, Thermo Fisher Scientific, Waltham, MA, USA), with yields ranging from 2.1 to 250 ng/µL. RNA integrity was assessed using the Agilent RNA ScreenTape Assay (RNA ScreenTape Analysis, #5067-5576, #5067-5577, #5067-5578, Agilent, Santa Clara, CA, USA) on the 4200 TapeStation system (4200 TapeStation System, #G2991BA, Agilent, Santa Clara, CA, USA) and analyzed with TapeStation Analysis Software (Agilent Tapestation Software, Agilent, Santa Clara, CA, USA). RNA integrity number (RIN) values ranged from 1.5 to 3.2, while DV200 values varied from 12.5% to 59.1%. FFPE RNA library preparation was carried out using the Lexogen QuantSeq 3’ mRNA-Seq Library Prep Kit for Illumina (QuantSeq3’ mRNA-Seq Library Prep Kit for Illumina (FWD), #015, Lexogen GmbH, Wien, Austria), following the manufacturer’s instructions. Libraries were quantified using the dsDNA HS Assay Kit (Qubit dsDNA HS Assay Kit, #Q32854, Thermo Fisher Scientific, Waltham, MA, USA) and a Qubit 4 Fluorometer (Qubit 4 fluorometer, Thermo Fisher Scientific, Waltham, MA, USA). DNA integrity was assessed with the Agilent DNA ScreenTape Assay (Genomic DNA ScreenTape Analysis, #5067-5365, #5067-5366, Agilent, Santa Clara, CA, USA) on the 4200 TapeStation system (4200 TapeStation System, #G2991BA, Agilent, Santa Clara, CA, USA). DNA yields ranged from 0.154 to 0.570 ng/µL. Libraries were equimolarly pooled and sequenced on a NextSeq 500 platform (NextSeq 500 System, Illumina Inc, San Diego, CA, USA) to an average depth of 7 × 10⁶ single-end reads. FastQC method was perform a quality check of FASTQ files ^52^. Fastp and SortMeRNA, and STAR algorithms were used to pre-process and align reads to the reference genome GRCh38, respectively ^54,56^. RSEM method was used to generate raw count matrix with hs ensembl.h38.v112 annotation ^55^.

### Semi-quantitative assessment of lymphocytic infiltration

Lymphocytic infiltration was scored on the 34 available hematoxylin and eosin (H&E)–stained diagnostic FFPE biopsy sections etrieved from our Institute using a four-tier scale based on both density and distribution of lymphoid cells within the tumor stroma: weakly positive (+), moderately positive (+/++), positive (++), and strongly positive (+++). Attention was given to the presence of lymphoid aggregates or clusters, which were recorded when present. Semi-quantitative lymphocytic scores were correlated with clinical outcome during AS.

### Targeted transcriptional profiling of FFPE DF biopsies

Targeted gene expression analysis was performed using the following NanoString nCounter panels: PanCancer Immune Profiling Panel (nCounter Human PanCancer Immune Profiling Panel, Bruker Corporation, Bothell WA, USA) and Fibrosis Panel (nCounter Fibrosis Panel, Bruker Corporation, Bothell WA, USA). The experiment was performed following the manufacturer instructions. Briefly, probes (capture and reporter) and a total amount of 200 ng mRNA were hybridized overnight at 65 °C for 16 h. Samples were then scanned at maximum resolution capabilities (555 FOV) using the nCounter Digital Analyzer (Bruker Corporation, Bothell WA, USA). Quality control of samples, data normalization, and advanced data analysis were performed using nSolver software 4.0 (nSolver 4.0 Analysis Software, Bruker Corporation, Bothell WA, USA). Genes differentially expressed between DF from progressing patients and DF from regressing patients have been identified with DESeq2 R package ^57^. Tests for DEGs identification are adjusted by FDR.

### DNA extraction from whole blood and HLA typing

DNA required for HLA-Typing was extracted from whole blood using QIAamp DNA Blood Mini Kit (QIAamp DNA Blood Mini Kit, #51104, QIAGEN, Hilden, Germany), following the manufacturer instructions. The extracted DNA was quantified by NanoDrop 2000 spectrometer (NanoDrop™ 2000 Spectrophotometers, #ND-2000, Thermo Fisher Scientific, Waltham, MA, USA). The DNA yield ranged from 11 to 130 ng/µl.

HLA typing of locus HLA-DRB1/B3/B4/B5 was performed by using NGSgo-AmpX v2 HLA-DRB1 (NGSgo®-AmpX v2 HLA-DRB1 Whole Gene, #7370622, GenDX, Chicago, IL, USA) and NGSgo-AmpX v2 HLA-DRB3/B4/B5 kit (NGSgo®-AmpX v2 HLA-DRB3/4/5, # 7970762, GenDX, Chicago, IL, USA), following the manufacturer instructions. A sample of known typing was used as control. The quality of the amplified DNA was assessed by agarose gel run, the amplification was considered valid only if each amplicon had the correct locus weight. Quantification of DNA was performed by dsDNA BR Assay Kit (Qubit dsDNA BR Assay Kit, #Q32850, Thermo Fisher Scientific, Waltham, MA, USA) and Qubit 4 Fluorimeter (Qubit 4 fluorometer, Thermo Fisher Scientific, Waltham, MA, USA). The libraries were prepared using the NGSgo-LibrX kit (NGSgo®-LibrX, #2342605, GenDx, Chicago, IL, USA), following the manufacturer instructions. The libraries DNA was quantified by dsDNA BR Assay Kit (Qubit dsDNA BR Assay Kit, ##Q32850, Thermo Fisher Scientific, Waltham, MA, USA) and Qubit 4 Fluorimeter (Qubit 4 fluorometer, Thermo Fisher Scientific, Waltham, MA, USA) and properly diluted to reach a final concentration of 55-76 pM. NGS was performed by iSeq100 (iSeq100, #20021535, Illumina Inc, San Diego, CA, USA). Data were analyzed by NGSengine (NGSengine®, #4109940, GenDx, Chicago, IL, USA) to assign the specific HLA allele.

### Bioinformatic analyses

Transcriptomic profile matrix was generated by RNA-Seq experiments. In the preprocessing step, genes with no count or low variability among all samples were filtered out, by using *filterByExpr* function implemented into the *edgeR* package ^58^. At the same time, genes without an official symbol were discarded, while those with same symbol were collapsed by summing their counts. Filtered data were normalized using the *Trimmed Mean of M-values* (TMM), implemented into the *edgeR* package. Samples with a third-quartile expression exceeding the maximum first quartile across samples were considered outliers and excluded. Differential expression analysis was performed by removing data heteroscedasticity with the voom method implemented into the edgeR package, and employing a linear model to evaluate transcriptomic changes using the limma package ^59^. For pre-ranked Gene Set Enrichment Analysis (GSEA) ^60^ the t-statistic was used on *hallmark* (H) and *immunologic signature* (C7) collections of the *Molecular Signature Database* (MSigDB) and on literature-derived or custom signatures (**Suppl. Table S3**), by using the fgsea package ^61^. An FDR threshold of 0.05 was applied to assess statistical significance pathways. Enrichment map was generated using enrichplot R package (v1.24.4) ^62^ with additional filtering and clustering optimization steps. Enriched pathways similarity was calculated using the Jaccard coefficient (JC) based on overlapping leading-edge genes. Each node corresponds to a significantly enriched pathway (p_adj_<0.05), where node size is proportional to the number of leading-edge genes and node color reflects the normalized enrichment score (NES). Edge thickness represents the JC. Edges were filtered using the minimum edge weight of 0.1, and isolated nodes were removed to improve readability. K-means clustering was used to group pathways, where the optimal number of clusters is selected by gap statistic and cluster labels were manually assigned based on the dominant biological themes within each cluster. The MCP-counter ^23^ deconvolution method was applied to estimate the abundance of immune cell subpopulations from bulk gene expression implemented into *immunedeconv* package.

Reactome pathway enrichment was performed using the ReactomePA package ^63^; pathways with FDR-adjusted p-value < 0.10 were considered statistically significant.

Protein expression differences at baseline were evaluated using the *Olink Analyze* package ^64^; proteins with FDR < 0.10 were retained and ranked by estimated log2 fold change (log2fc).

All transcriptomic and proteomic analyses were conducted in R (version 4.4.1).

### Statistical analyses

Descriptive statistics were used to summarize patients’ characteristics. Numerical variables were reported as medians with interquartile ranges (IQR), while categorical variables were described using absolute and relative frequencies.

Two time-to-event endpoints were evaluated: progression-free survival (PFS), defined as the time from under AS to the first documented disease progression, and treatment-free survival (TFS), defined as the time from study entry to initiation of antitumor treatment; observation times for patients who did not experience progression (for PFS) or did not initiate treatment (for TFS) were censored to the date of last follow-up.

Kaplan-Meier curves were represented over a 24-month time to describe PFS and TFS according to baseline covariates. Selected covariates were recategorized as follows: *CTNNB1* mutation status was grouped into three categories, WT/S45P/Other (reference; including wild-type, S45P, and other/rare variants), S45F, and T41A; anatomic site was classified as abdominal wall (reference) versus other sites (trunk, extremity, head and neck, and intra-abdominal); age was dichotomized at the sample median. Univariate Cox models were fitted to estimate hazard ratios (HRs) with 95% confidence intervals (CIs) for each covariate.

Multivariate Cox proportional hazards models were fitted separately for PFS and TFS to evaluate the independent association of baseline covariates, age, tumor size, *CTNNB1* mutation status, anatomic site, and autoimmune disease status, with each endpoint. Age and tumor size were analyzed as continuous variables and modeled using restricted cubic splines with three knots placed at empirical quartiles to accommodate for potential nonlinear associations. A two-sided significance level of 0.05 was used for all comparisons.

Statistical Analyses were performed with R version 4.4.1.

## Availability of data and materials

The data presented in the study are deposited in the GEO repository, accession numbers GSE324464 and GSE324484.

## Acknowledgements

This work was supported by Italian Ministry of Health through GR-2016-02362609 to C.C. and V.V., by 5xmille funds for healthcare research (Ministry of Health) to C.C. and L.B., “Ricerca Corrente” to Sa.P. St. P. was supported by “Institutional 5x1000 Funds BRI 2021”.

## Author contributions

M.F., E.P., A.G., V.V., C.C., L.R., and Sa.P. conceived and designed the study. L.B., F.R., L.D.C., A.L., M.Z., F.P., P.C., B.V., and C.B. were responsible for data acquisition. L.B., St.P., Y.Z., G.T., R.M., J.G., A.M., BE.L., V.V., and C.C. performed data analysis and interpreted the results. All authors contributed to writing the manuscript, critically revised its content, and approved the final version.

## Supplementary Information

**Supplementary Figures 1.** Gating strategy used to analyze data from WB samples tested with Granulocytes WB panel. WB: whole blood.

**Supplementary Figures 2.** Gating strategy used to analyze data from fresh PBMC samples tested with Granulocytes PBMC panel. PBMC: Peripheral Blood Mononuclear Cells.

**Supplementary Figures 3.** Gating strategy used to analyze data from fresh PBMC samples tested with Monocytes PBMC panel. PBMC: Peripheral Blood Mononuclear Cells.

**Supplementary Figures 4.** Gating strategy used to analyze data from frozen PBMC samples tested with T cell exhaustion /activation /Treg PBMC panel. Treg: T regulatory cells. PBMC: Peripheral Blood Mononuclear Cells.

**Supplementary Figures 5.** Gating strategy used to analyze data from frozen PBMC samples tested with T helper PBMC panel. PBMC: Peripheral Blood Mononuclear Cells.

**Supplementary Figures 6.** Gating strategy used to analyze data from surgical sample cell suspension samples tested with Immune infiltrate-tumor cell suspension panel.

**Supplementary Figure 7. Kaplan-Meier curves for PFS and TFS according to clinical variables.** (A-E) Progression-free survival (PFS) stratified by age, sex, β-catenin mutation status, tumor site and autoimmune disease. (F-J) Treatment-free survival (TFS) stratified by the same variables. For each comparison, hazard ratios (HR), 95% confidence intervals (CI), and corresponding p-values are reported. P values < 0.05 were considered statistically significant.

**Supplementary Figure 8.** Blood data of reclassified prog and reg patients.

**Supplementary Figure 9.** Proteomic blood profiling of prog and reg patients.

**Supplementary Table S1.** Flow cytometry antibody panels.

**Supplementary Table S2.** Flow cytometry antibody list.

**Supplementary Table S3.** List of custom immune gene sets.

**Supplementary Table S4.** Detailed patient clinical and tumor characteristics and samples analyzed in the study. The different analyses performed are reported.

**Supplementary Table S5.** Flow Cytometry data analysis PROG vs REG. Differences with p-value < 0.05 were considered significant.

**Supplementary Table S6.** Flow cytometry variables resulted significant by univariate analysis with logistic regression.

**Supplementary Table S7.** Reactome analysis of blood proteomic data. Pathways with p.adjust < 0.05 were considered significant.

**Supplementary Table S8.** Immune-related gene sets significantly enriched in whole blood from DF patients with regression compared to those with progression during AS (p_adj_ < 0.05).

**Supplementary Table S9.** Immune-related gene sets significantly enriched in whole blood from DF patients reclassified as regressors (rREG) compared to progressors (rPROG) during AS (p_adj_ < 0.05). Gray shading indicates gene sets also significant in the corresponding RECIST-based analysis.

**Supplementary Table S10.** Gene sets significantly enriched in biopsies from DF patients with regression compared to those with progression during AS (p_adj_ < 0.05).

**Supplementary Table S11.** Immune-related gene sets significantly enriched in biopsies from DF patients reclassified as regressors (rREG) compared to progressors (rPROG) during AS (p_adj_ < 0.05). Gray shading indicates gene sets also significant in the corresponding RECIST-based analysis.

**Supplementary Table S12.** HLA-DRB1/3/4/5 Genotyping Results.

**Supplementary Table S13.** Gene sets significantly enriched in biopsies from DF patients with autoimmune disease compared to those without autoimmune disease (p_adj_ < 0.05).

