## Supplemental Figures 1-9 for "Active Surveillance Reveals a Systemic Pro-Resolving Th2 Immune Program Linked to Desmoid Tumor Regression"

Supplementary Figure S1

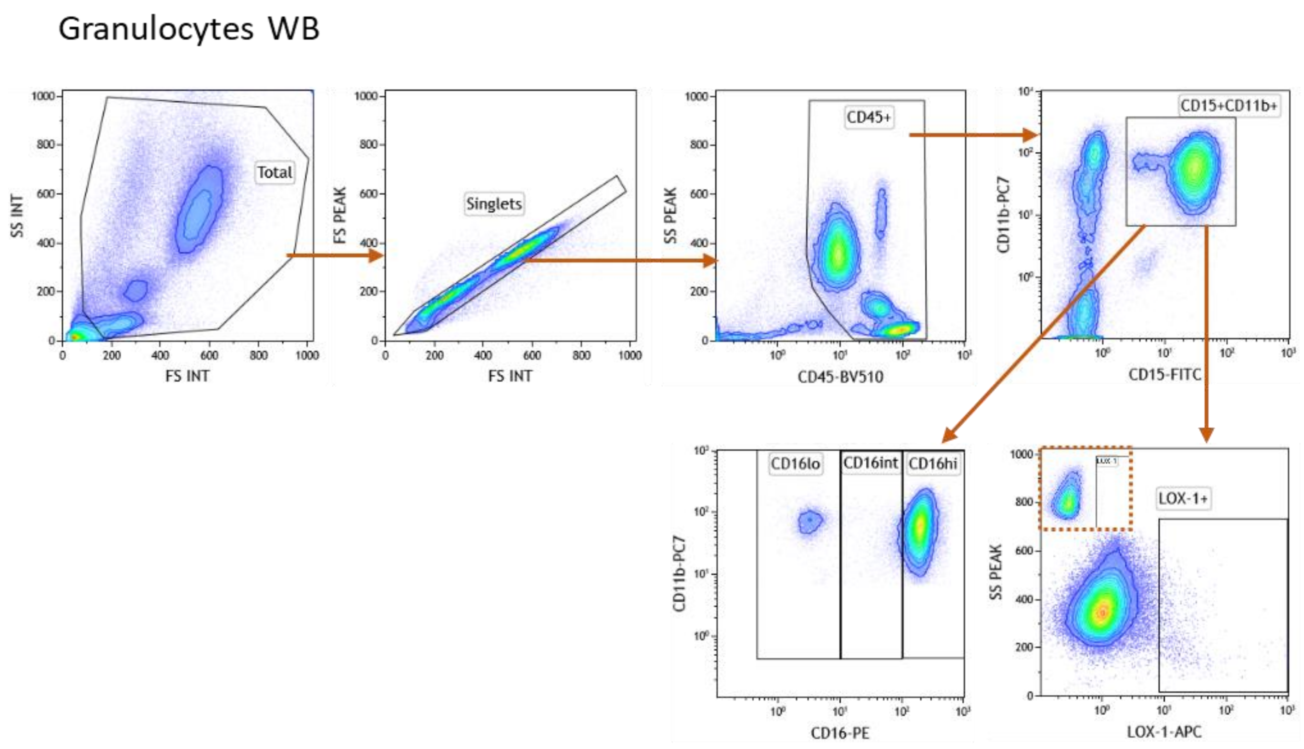


**Supplementary Figure S1:** Gating strategy used to analyse data from WB samples tested with Granulocytes WB panel.

### Supplementary Figure S2

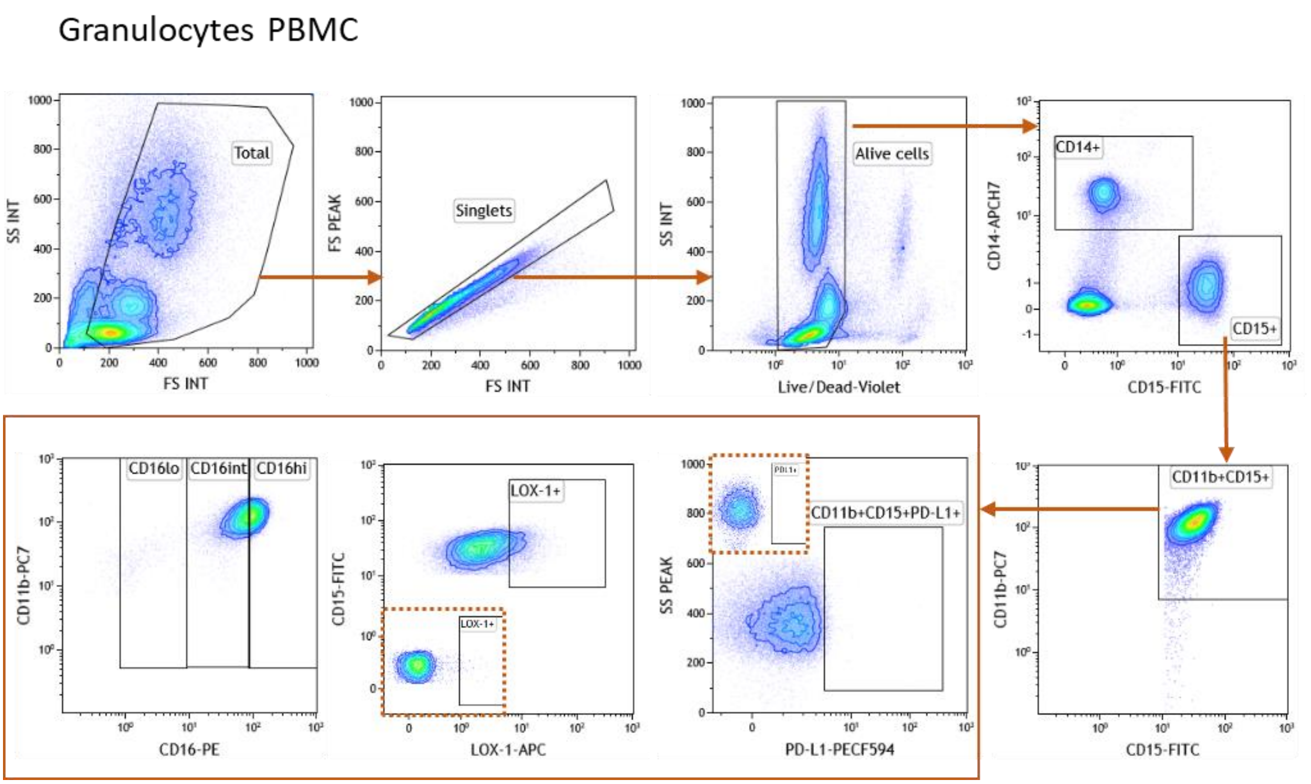


**Sipplementary Figure S2:** Gating strategy used to analyse data from fresh PBMC samples tested with Granulocytes PBMC panel.

### Supplementary Figure S3

#### Monocytes PBMC

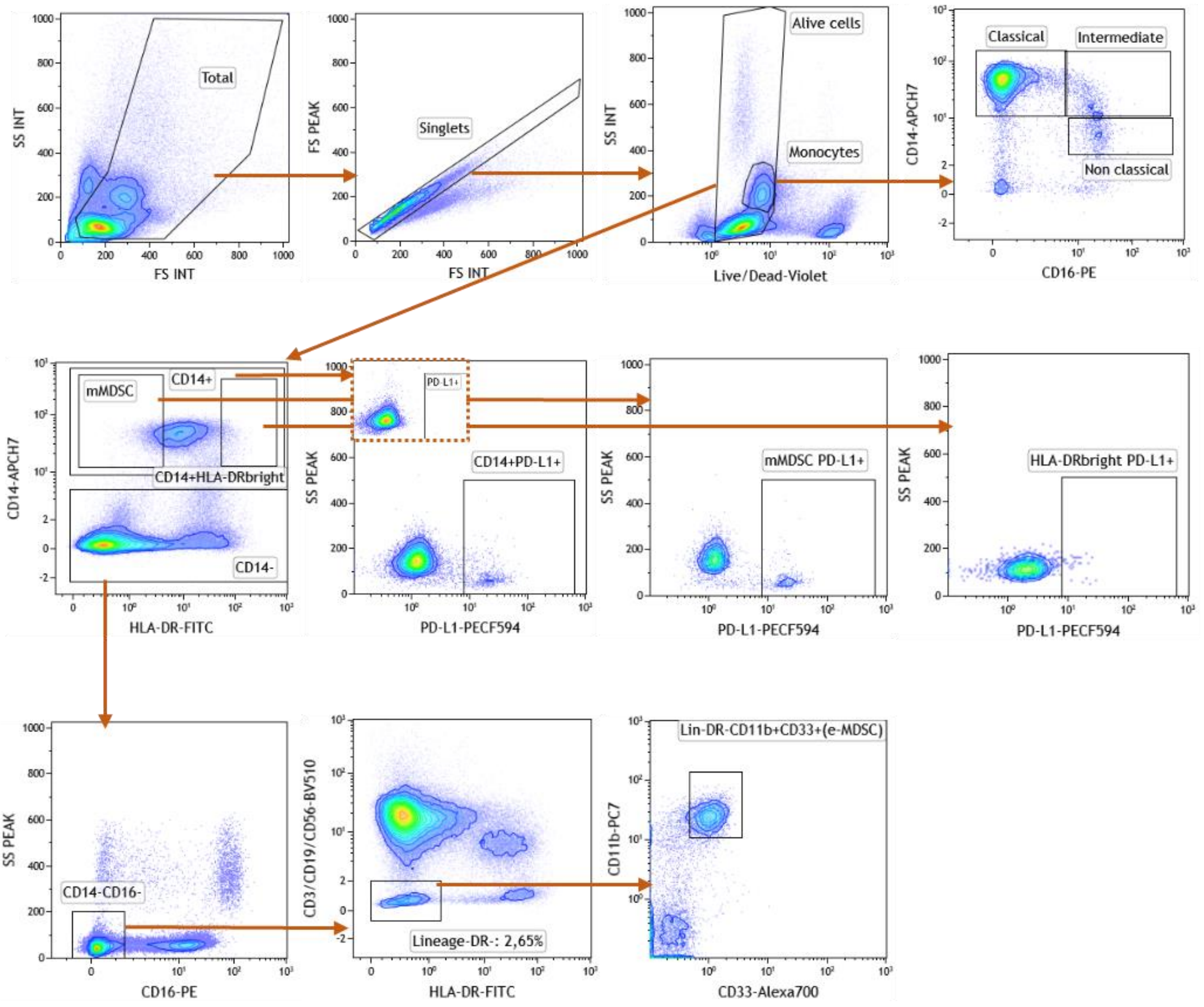


**Supplementary Figure S3:** Gating strategy used to analyse data from fresh PBMC samples tested with Monocytes PBMC panel.

### Supplementary Figure S4

#### T cell exhaustion/activation/Treg PBMC

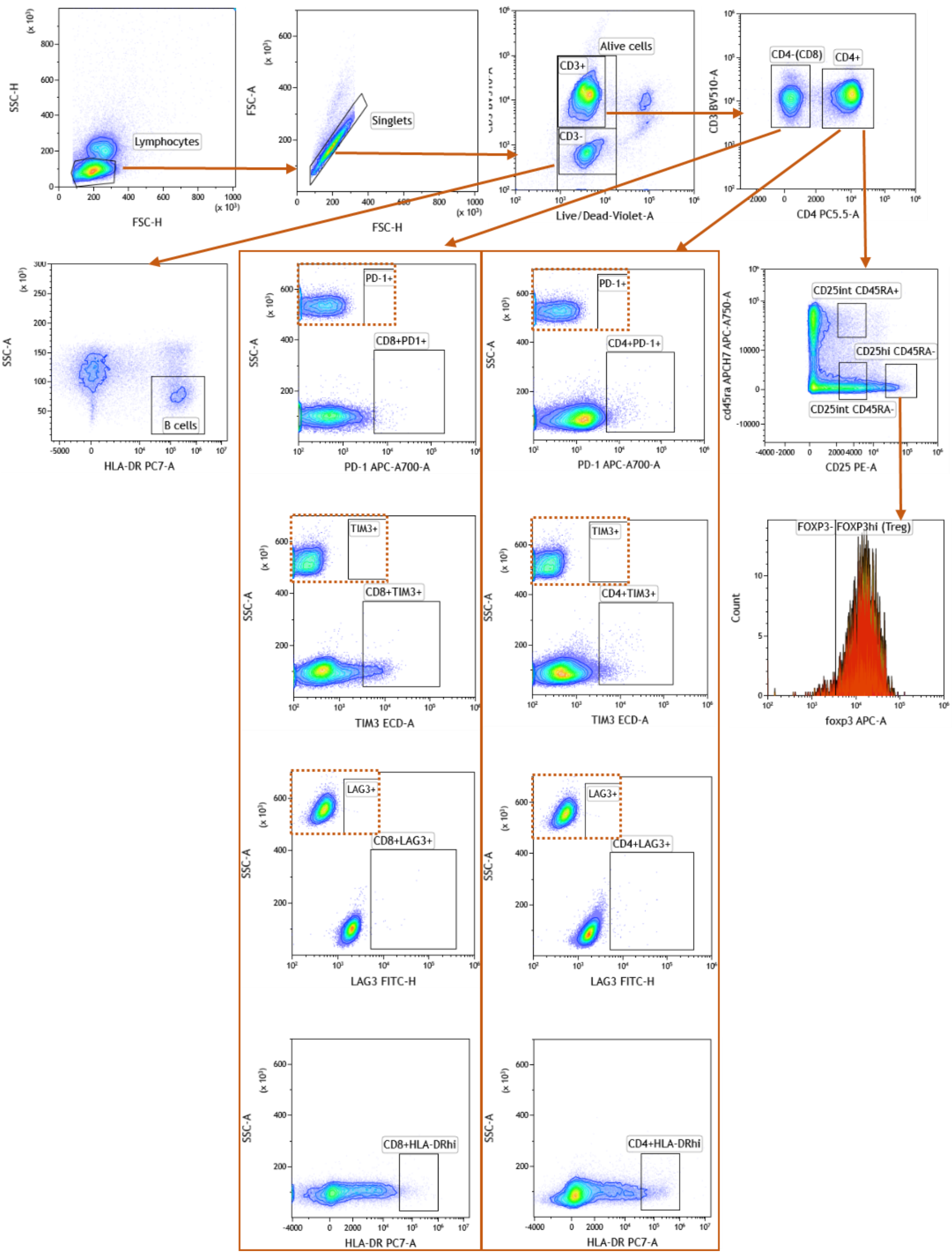


**Supplementary Figure S4:** Gating strategy used to analyse data from frozen PBMC samples tested with T cell exhaustion /activation /Treg PBMC panel.

### Supplementary Figure S5

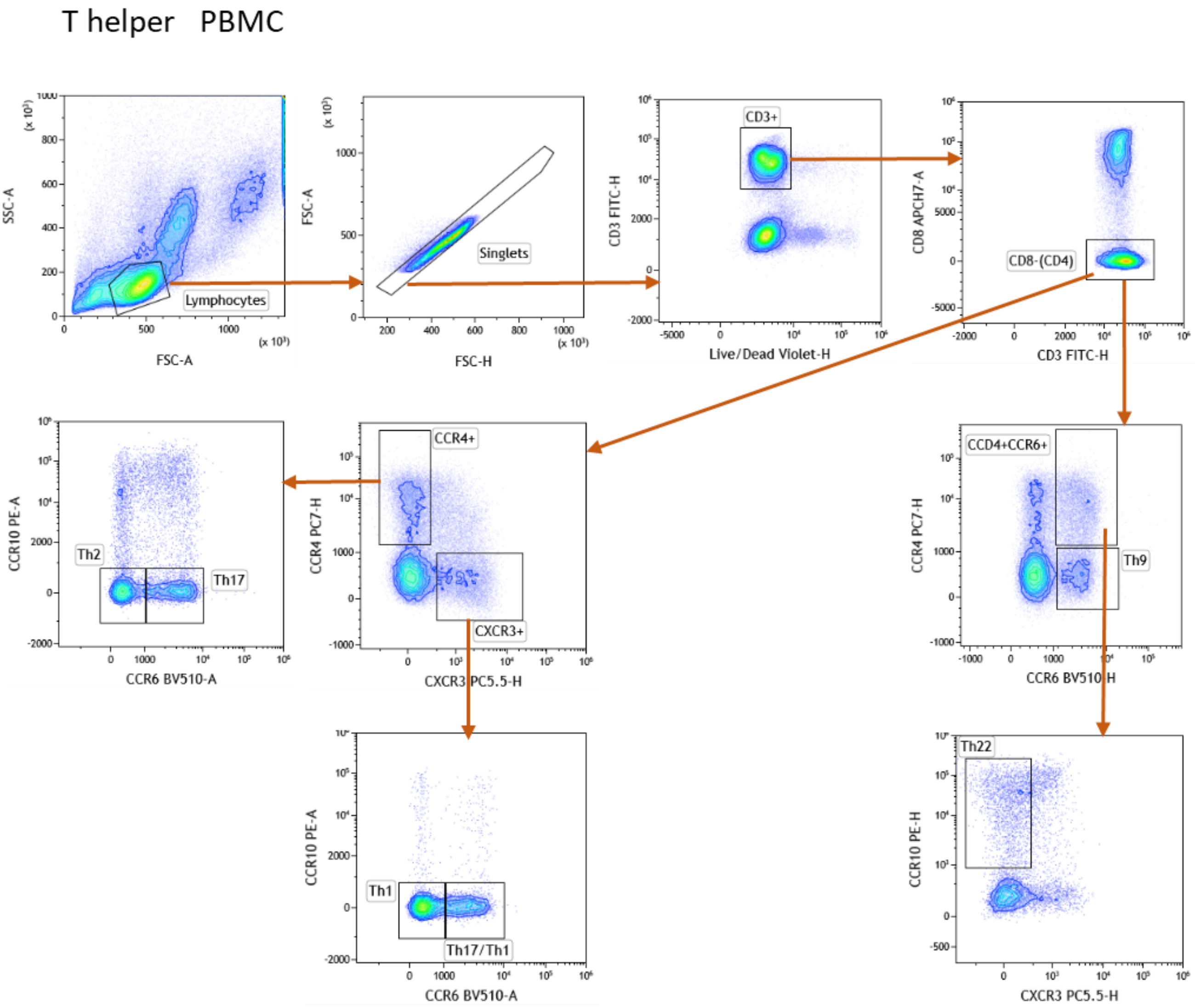


**Supplementary Figure S5:** Gating strategy used to analyse data from frozen PBMC samples tested with T helper PBMC panel.

Supplementary Figure S6

Immune infiltrate tumor cell suspension

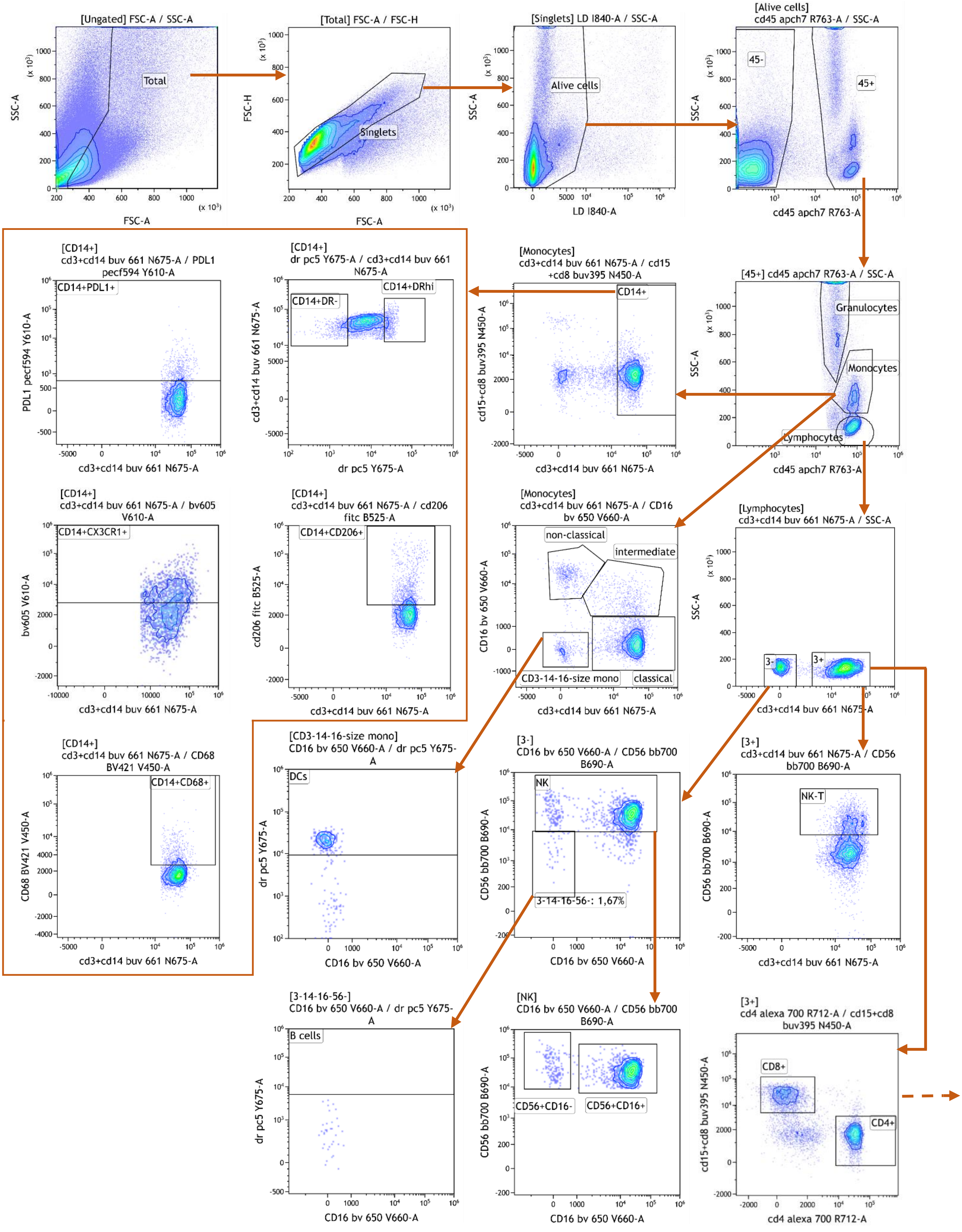

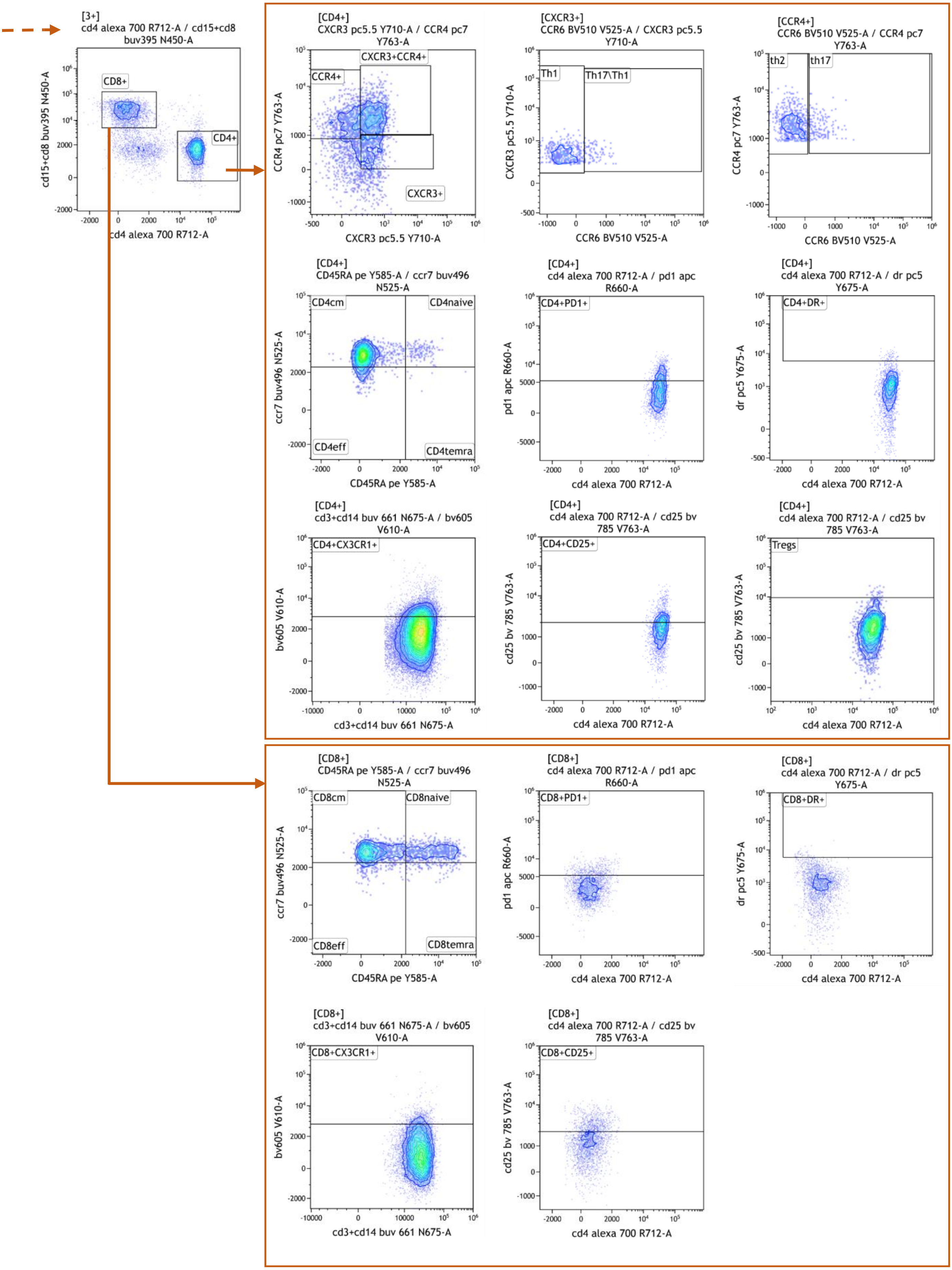


**Supplementary Figure S6:** Gating strategy used to analyse data from surgical sample cell suspension samples tested with Immune infiltrate-tumor cell suspension panel.

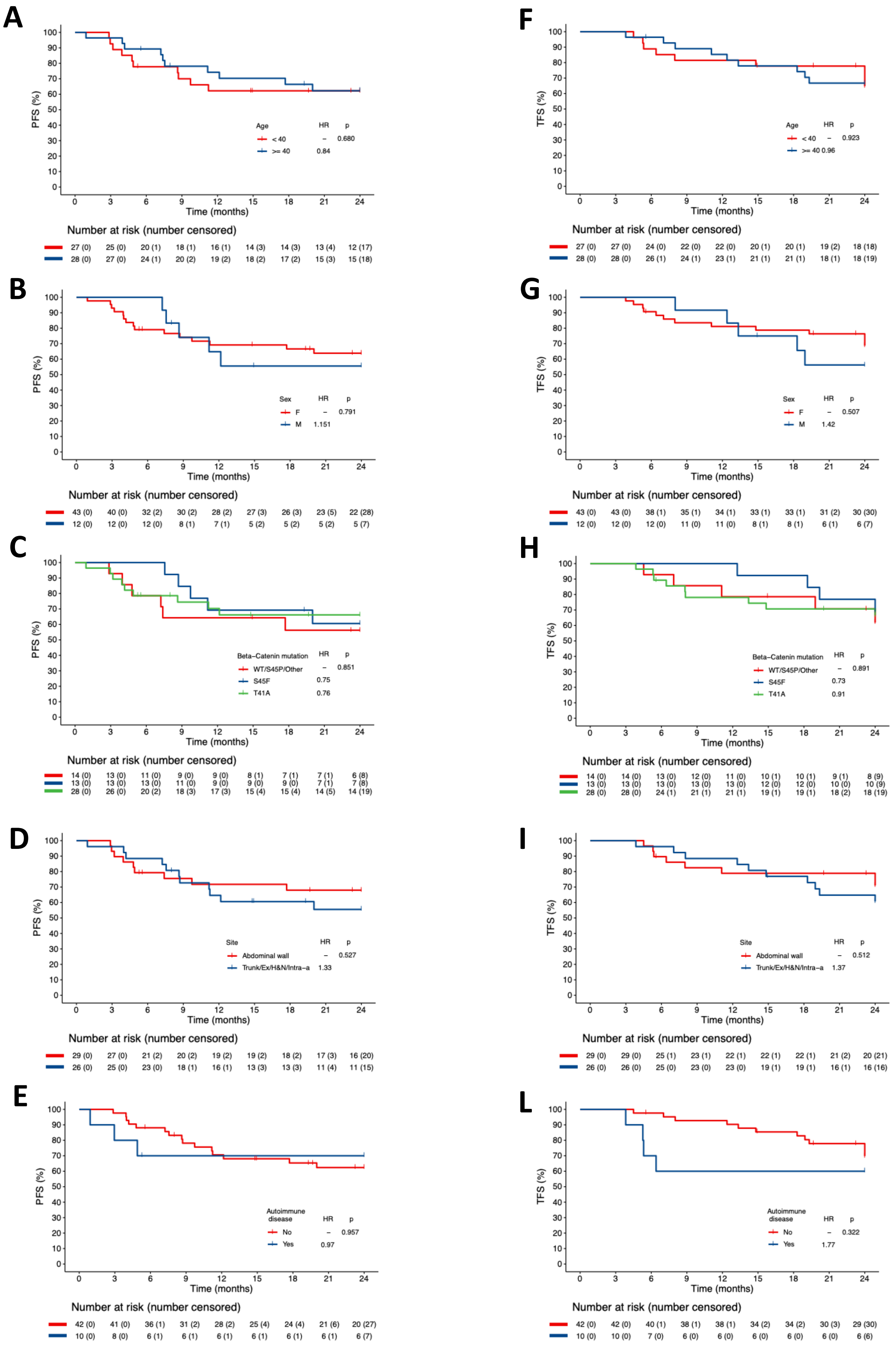


**Supplementary Figure S7**

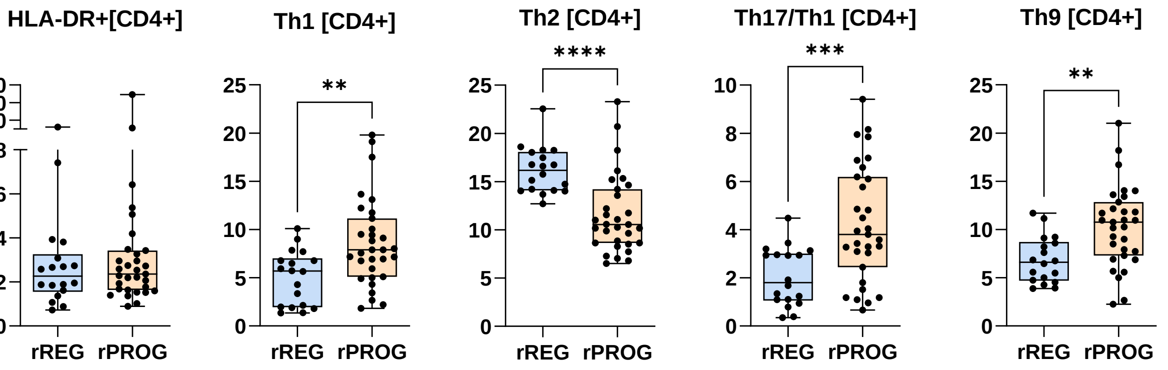


**Supplementary Figure S8**

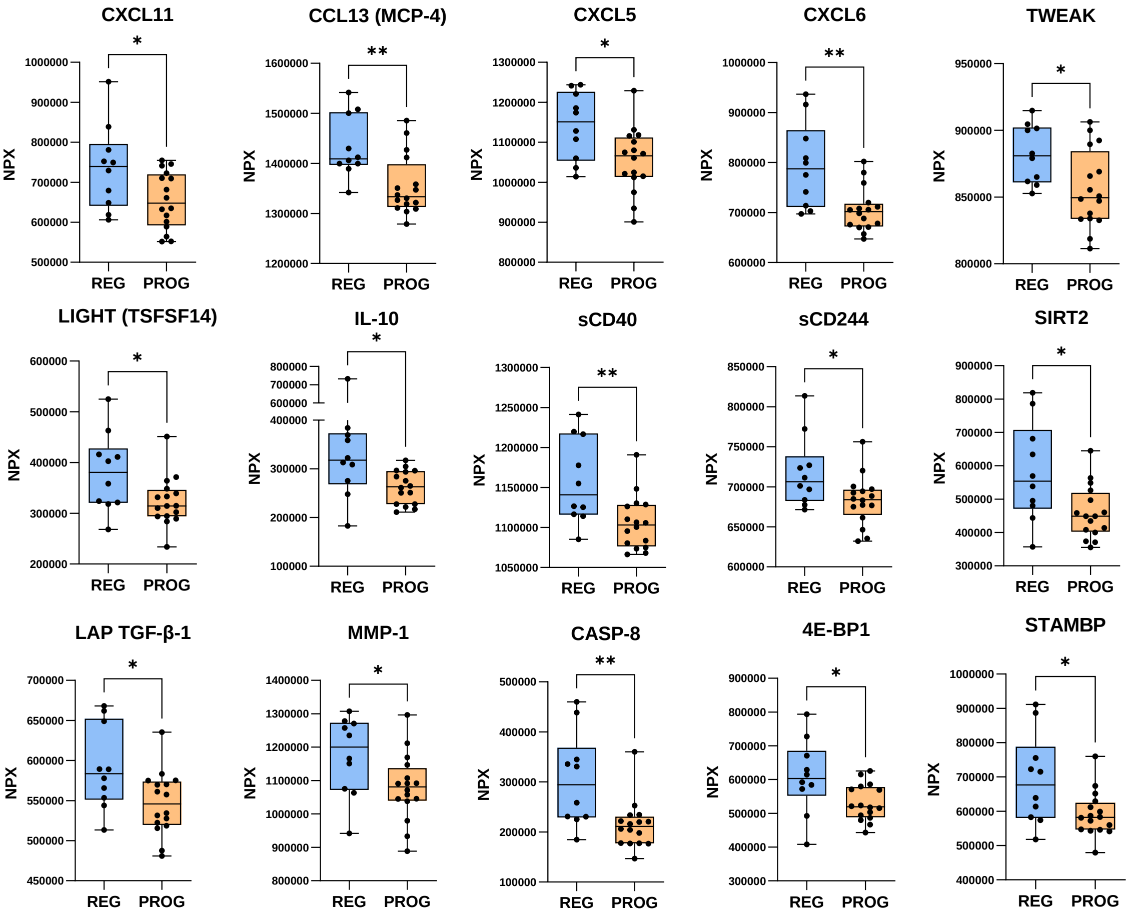


### Supplementary Figure S9
